# Autoantibody and hormone activation of the thyrotropin G protein-coupled receptor

**DOI:** 10.1101/2022.01.06.475289

**Authors:** Bryan Faust, Isha Singh, Kaihua Zhang, Nicholas Hoppe, Antonio F. M. Pinto, Yagmur Muftuoglu, Christian B. Billesbølle, Alan Saghatelian, Yifan Cheng, Aashish Manglik

**Affiliations:** Department of Pharmaceutical Chemistry, University of California, San Francisco, CA, USA; Department of Biochemistry & Biophysics, University of California, San Francisco, CA, USA; Biophysics Graduate Program, University of California, San Francisco, CA, USA; Peptide Biology Lab and Mass Spectrometry Core, Salk Institute for Biological Studies, La Jolla, CA, USA; Stanford University School of Medicine, Stanford, CA, USA; Howard Hughes Medical Institute, University of California, San Francisco, CA, USA; Department of Anesthesia and Perioperative Care, University of California, San Francisco, CA, USA

**Keywords:** G protein-coupled receptor, autoantibody, autoimmune disease, thyroid hormone

## Abstract

Thyroid hormones are vital to growth and metabolism. Thyroid hormone synthesis is controlled by thyrotropin (TSH), which acts at the thyrotropin receptor (TSHR). Autoantibodies that activate the TSHR pathologically increase thyroid hormones in Graves’ disease. How autoantibodies mimic TSH function remains unclear. We determined cryogenic-electron microscopy structures of active and inactive TSHR. In inactive TSHR, the extracellular domain lies close to the membrane bilayer. TSH selects an upright conformation of the extracellular domain due to steric clashes between a conserved hormone glycan and the membrane bilayer. An activating autoantibody selects a similar upright conformation of the extracellular domain. Conformational changes in the extracellular domain are transduced to the seven transmembrane domain via a conserved hinge domain, a tethered peptide agonist, and a phospholipid that binds within the seven transmembrane domain. Rotation of the TSHR ECD relative to the membrane bilayer is sufficient for receptor activation, revealing a shared mechanism for other glycoprotein hormone receptors that may also extend to G protein-coupled receptors with large extracellular domains.

## Introduction

The thyroid gland regulates organ development and homeostatic metabolism in all vertebrates via the thyroid hormones triiodothyronine (T_3_) and thyroxine (T_4_)^1^. Synthesis and secretion of the thyroid hormones is controlled by a hypothalamic-pituitary-thyroid homeostatic signaling axis^2^. Hypothalamic sensing of low thyroid hormone levels induces secretion of the pituitary thyrotropin hormone (also called thyroid stimulating hormone, TSH), which acts at the thyrotropin receptor (TSHR), a G protein-coupled receptor (GPCR) located on thyroid follicles^3^. Activation of heterotrimeric G_s_ and G_q_ signaling pathways downstream of TSHR leads to thyroid hormone production, closing the negative feedback loop to set physiological thyroid hormone levels^4^.

Dysregulation of the central hypothalamic-pituitary-thyroid signaling axis leads to inappropriately increased or decreased thyroid hormone levels, leading to a disease burden that affects approximately 5% of the world population^5^. Hypothyroidism predominantly stems from iodine deficiency or autoimmune inflammation of the thyroid. The primary cause of hyperthyroidism in countries without iodine deficiency is Graves’ disease, an autoimmune disorder leading to inappropriate activation of the TSHR by autoantibody thyroid stimulating immunoglobulins (TSI)^6,7^. Direct activation of the TSHR by TSI overcomes the physiological hypothalamic-pituitary-thyroid negative feedback loop, leading to elevated thyroid hormones despite low levels of serum TSH. Despite the central role of the TSHR in regulating thyroid hormone physiology, no currently approved medications directly target this receptor to treat thyroid diseases^8^. Current interventions either target thyroid hormone synthesis, or in more severe cases, destroy the thyroid gland leading to a lifelong need for thyroid hormone replacement therapy^9^.

A deeper understanding of how TSH physiologically activates the TSHR and how TSI pathologically stimulate the TSHR would enable approaches to precisely tune the hypothalamic-pituitary-thyroid signaling axis to correct thyroid disease. A central challenge, however, has been the design of TSHR selective molecules, due in part to a limited understanding of the structure and dynamics of TSHR function. The TSHR shares significant similarity to two homologous receptors that are critically important in reproductive physiology: the follicle-stimulating hormone receptor (FSHR) and the luteinizing hormone– choriogonadotropin receptor (LH/CGR). While the FSHR is specific for the follicle stimulating hormone (FSH), the LH/CGR is activated by both luteinizing hormone (LH) and chorionic gonadotropin (CG). A central hallmark of TSH, FSH, LH and CG, together termed glycoprotein hormones, is their complex glycosylation required for biological activity^10–12^. Structures of the FSHR and LH/CGR bound to FSH and CG, respectively, have illuminated key aspects of glycoprotein hormone recognition and receptor activation. However, these studies leave unanswered how glycosylation drives hormone activity, and for the TSHR, how pathogenic autoantibodies mimic TSH action. Here, we use a combination of cryogenic electron-microscopy (cryo-EM) and signaling studies to illuminate the molecular basis of action of TSH and autoantibodies at the TSHR. Our studies provide a model for physiological and pathological activation of the TSHR and a general activation mechanism for the glycoprotein hormone receptor family.

## Results

### Structure of TSH-activated TSHR bound to G_s_

We first obtained a structure of activated TSHR bound to native human TSH (Fig. 1a). TSHR is naturally proteolytically cleaved within the extracellular domain leading to removal of residues 317 to 366, a region termed the “C-peptide”^13^. While the physiological relevance of C-peptide excision remains unclear, previous biochemical studies have demonstrated that cleavage in this domain leads to receptor instability^14^. To simplify purification of intact receptors for structural studies, we generated a TSHR construct devoid of the C-peptide, which could be purified as a single intact receptor construct. In agreement with prior studies demonstrating that this region is not required for proper receptor folding and signaling^14^, removal of the C-peptide did not affect TSH or TSI activation of G_s_ signaling (Supplementary Fig. 1). To further improve the expression and purification of active TSHR, we generated a construct with a C-terminal fusion of an engineered “mini” Gα_s_ protein. This miniGα_s_ protein contains only the Ras-like GTPase domain of Gα_s_ and is thermostabilized to interact with a GPCR in a nucleotide-independent manner^15^. We purified this construct, termed TSHR-miniG_s_, in complex with native TSH purified from human pituitary, recombinantly expressed Gβ_1_γ_2_, and the G_s_ stabilizing nanobody 35 (Nb35)^16^. The resulting preparation of TSH-activated, G_s_-bound TSHR was stable enough to enable single particle cryo-EM. Because of conformational heterogeneity caused by flexibility between the TSHR extracellular domain (ECD) and the seven transmembrane domain (7TM) bound to G_s_, we separately classified and refined these two regions of the complex to yield a 3.4 Å resolution map of the TSHR ECD bound to TSH and a 2.9 Å map of the TSHR 7TM bound to the G_s_ heterotrimer (Supplementary Fig. 2). A composite map from these individual reconstructions enabled us to build an atomic model for the majority of TSHR and the G_s_ heterotrimer, and key interacting regions between TSH and TSHR (Fig. 1b).

**Fig. 1.**
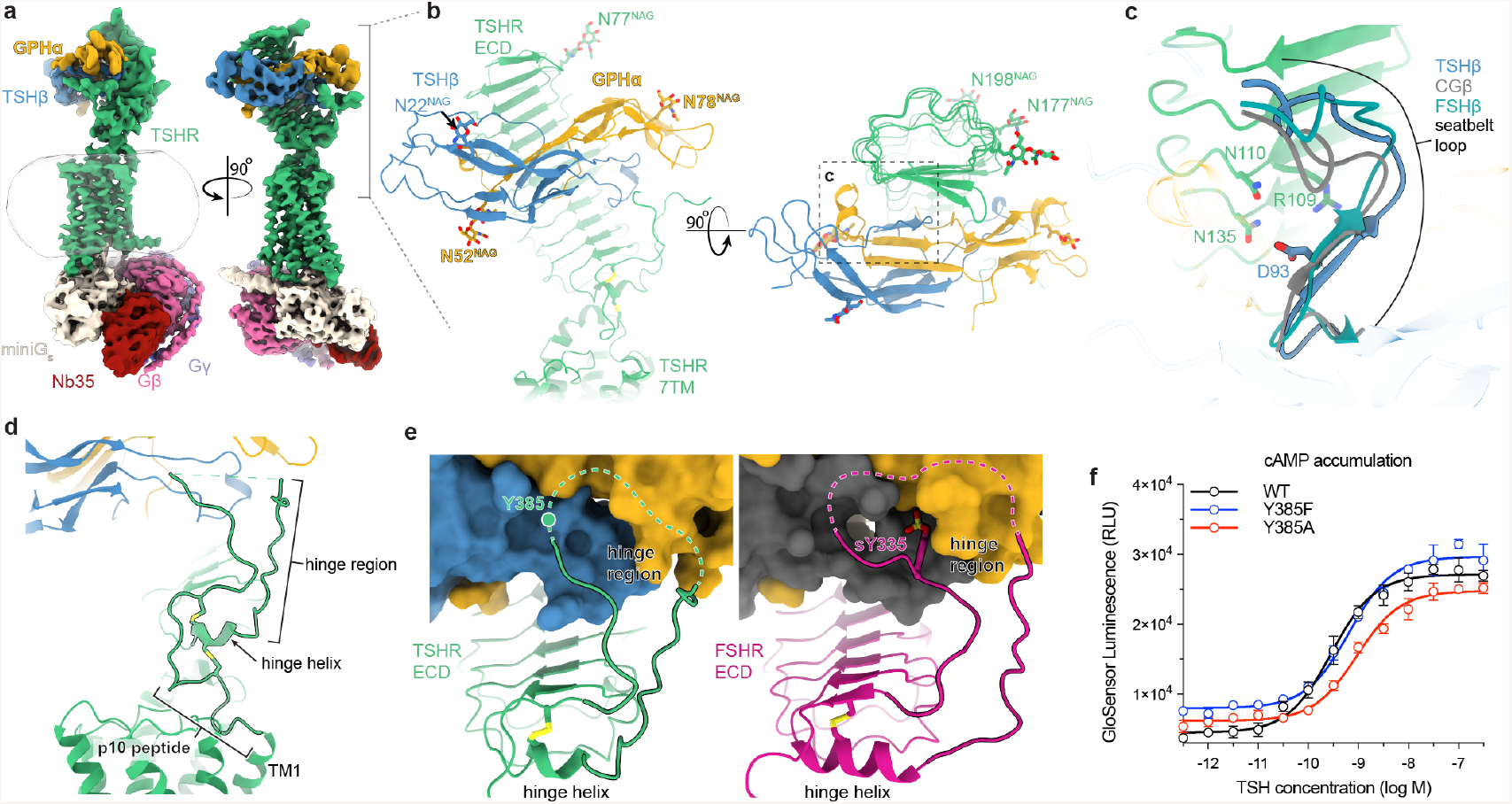
Cryo-EM structure of TSHR bound to native human TSH. **a)** Cryo-EM map of TSH-TSHR-G_s_-Nb35 complex. **b)** TSH binds to the TSHR extracellular domain (ECD). Resolved N-linked glycans are highlighted for both TSH and TSHR (N-acetylglucosamine, NAG). **c)** Close-up view of the seatbelt loop region. D93 anchors the seatbelt loop in TSHβ to the TSHR ECD. The other regions of the seatbelt loop differ in backbone conformation among glycoprotein hormones bound to their cognate receptors. **d)** The TSHR ECD connects to the 7TM domain via a hinge domain, comprised of a hinge region and a hinge helix, and a conserved p10 peptide. **e)** Comparison of hinge region interactions between TSH-bound TSHR and FSH-bound FSHR (PDB ID: 4AY9). The sulfotyrosine residue Y335 (sY335) in FSHR binds in a cavity within FSH. In TSHR, the corresponding tyrosine residue (Y335) is in a disordered portion of the hinge region. **f)** Mutations to Y385 that prevent tyrosine sulfation show similar TSH-dependent activity at TSHR. Signaling data points represent the global fit of grouped triplicate measurements at each concentration ±SD from 2 independent experiments.

TSH binds to a concave surface of the TSHR ECD leucine rich repeat (LRR) similar to previous structures of follicle stimulating hormone (FSH) bound to FSHR and chorionic gonadotropin (CG) bound to LH/CGR^17–19^ (Supplementary Fig. 3). The glycoprotein hormones are comprised of a common glycoprotein α chain (GPHα) and hormone-specific β chains. As previously observed for FSH and CG, the common GPHα chain of TSH makes two key interactions with the TSHR ECD. The C-terminus of the GPHα chain contacts the concave surface of the TSHR ECD, mediated primarily by hydrophobic interactions (Fig. 1b and Supplementary Fig. 3). A short alpha helix, comprised of residues 40-48, contacts the lateral edge of the TSHR ECD (Fig. 1b and Supplementary Fig. 3). Both interactions are similar to previous structures of glycoprotein hormones bound to their cognate receptors^18,19^, supporting the importance of the common GPHα in hormone binding (Supplementary Fig. 3). Importantly, our structure resolves glycosylation at Asn52 of the GPHα chain (Fig. 1b), a modification required for glycoprotein hormone signaling^11,20^.

Selective binding of glycoprotein hormones to their receptors is determined by the hormone-specific β chains^21,22^. The primary interaction between the TSH-specific β chain (TSHβ) and the TSHR is via a 16-residue segment previously termed the “seatbelt” loop^23,24^. The activity of chimeric glycoproteins demonstrated that this region confers specificity within the glycoprotein hormone family^25,26^. TSHβ D93 anchors the seat-belt with the TSHR ECD. A similar aspartic acid in FSHβ and CGβ anchor these hormones to their cognate receptors, which supports the importance of this interaction in binding of all three glycoprotein hormones^27^ (Fig. 1c and Supplementary Fig. 3). While our structure does not resolve specific interactions that distinguish TSH binding to the TSHR, the backbone conformation of the seat-belt loop is different between TSH, CG, and FSH bound to their cognate receptors, suggesting that conformational differences in this region may underlie the molecular basis of glycoprotein hormone selectivity (Fig. 1c).

The TSHR ECD connects to the 7TM domain in a network of interactions between two regions that are critical to glycoprotein hormone receptor activation: the “hinge domain” and the “p10 peptide”^28,29,30^ (Fig. 1d). The hinge domain is further separated into a “hinge helix” at the base of the TSHR ECD and a more unstructured “hinge region”. The C-terminus of the hinge region connects to transmembrane helix 1 (TM1) of TSHR via the p10 peptide, a 10-amino acid link region that has been proposed to be a tethered intramolecular agonist for glycoprotein hormone receptors^30^. The hinge helix acts as a central hub to connect TSHR ECD, the hinge region, and the p10 peptide regions via two disulfide bonds.

Our structure resolves a portion of the hinge region backbone (Fig. 1d). Within the hinge region, previous studies have identified Tyr385 as an important residue in TSH-mediated signaling^31^. Sulfation of Tyr385 in TSHR, and a homologous tyrosine in FSHR and LH/CGR, has been proposed to be critical for glycoprotein hormone action^18^. Although a previous structure supported a direct recognition of sulfated tyrosine in the hinge region of FSHR in a cationic pocket on FSH, the electron density in this X-ray structure is poorly resolved in this region^18^. Our reconstruction of TSH bound to the TSHR revealed a distinct conformation of the hinge region when compared to the structure of FSH bound to FSHR (Fig. 1e), with no clear density for binding of a sulfated Tyr385 to TSH. Contrary to previous reports^31,32^, we found Y385 mutations (Y385F and Y385A) that would preclude sulfation retained similar TSH potencies and efficacies as the wild-type protein (Fig. 1f and Supplementary Fig. 1). The structure of TSH bound to active TSHR therefore revealed both similarities and differences in hormone recognition within the glycoprotein hormone receptor family.

### Conformational changes in TSHR activation

To understand how TSH activates the TSHR, we next determined an inactive-state structure of the receptor (Fig. 2a,b). We initially attempted to capture the inactive state of TSHR stabilized by the small molecule negative allosteric modulator Org 274179-0, which is proposed to bind within the 7TM domain of the TSHR^33^. Despite clear 2D class averages supporting a TSHR-micelle complex, our efforts to obtain a high-resolution reconstruction were unsuccessful (Supplementary Fig. 4). We speculated that this result likely was due to conformational flexibility of the TSHR ECD compared to the 7TM domain associated with constitutive activity of the TSHR^34^. To further stabilize an inactive conformation, we used a Fab fragment of the inverse agonist CS-17 antibody that suppresses constitutive activity of the TSHR^35^. We obtained a reconstruction of the TSHR:Org 274179-0:CS-17 complex to a global resolution of 3.1 Å, with high resolution features for the TSHR ECD and CS-17 Fab and lower resolution reconstruction of the TSHR 7TM domain (Fig. 2a and Supplementary Fig. 5). While this map enabled an accurate model for the TSHR ECD and CS-17 Fab, the resolution in the TSHR transmembrane regions limited our ability to model side chains accurately or define high resolution interactions between Org 274179-0 and the 7TM domain.

**Figure 2.**
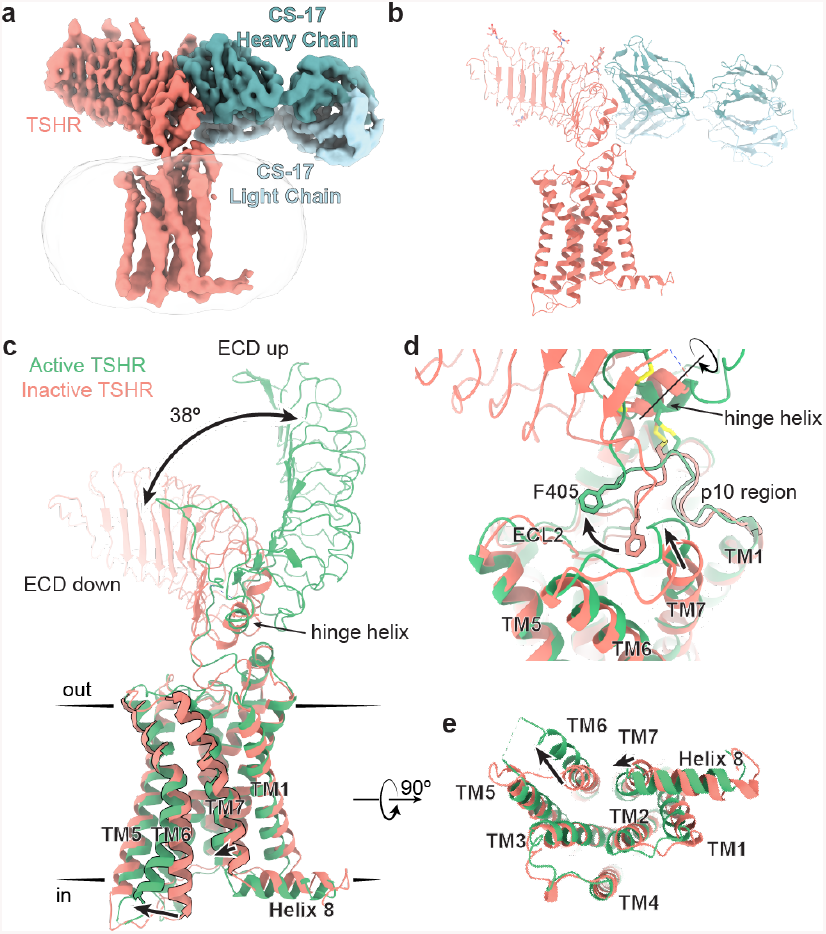
Activation mechanism of TSHR revealed by inactive structure bound to inverse agonist CS-17. Cryo-EM map (**a**) and model (**b**) of inactive TSHR bound to CS-17 Fab. **c)** Structural comparison of inactive and active TSHR with the 7TM domain aligned. In inactive TSHR, the extracellular domain (ECD) is in a down conformation close to the membrane bilayer. In active TSHR, the ECD is in an up conformation. The ECD rotates 38° between the up and down states. **d)** Rotation of the ECD leads to a rotation of the hinge helix. The p10 peptide wraps around the hinge like a pulley; ECD rotation thereby leads to an extracellular displacement of the p10 peptide upon TSHR activation. Notably the extracellular end of TM7 moves inward. **e)** In the active state, TM6 of TSHR displaces outward by 14 Å, while TM7 moves inward by 4 Å.

The TSHR ECD is rotated 38° towards the 7TM domain and the membrane bilayer in the inactive state when compared to the active state bound to TSH. We term this the “down” state. By contrast, we label the ECD in the “up” state as observed for active TSHR bound to TSH (Fig. 2c). Importantly, the structure of the LRR domain itself does not change between the inactive and active states of the TSHR (root mean squared deviation (RMSD) of 0.8 Å). The ECD therefore moves as a rigid body between the inactive and active states. Comparison of these new TSHR structures with previously determined structures of LH/CGR shows a similar relative orientation of the ECD in the inactive and active conformations (Supplementary Fig. 6). We surmise that this activation-associated conformational change in the ECD is common to the glycoprotein hormone receptor family and likely extends to the broader LRR-linked GPCR family.

The TSHR 7TM domain displays several classic hallmarks of GPCR activation. Compared to the inactive state, TM6 of active TSHR is displaced outward by 14 Å to accommodate the α5 helix of miniGα_s_ and TM7 moves ∼4 Å inward relative to the transmembrane core of the receptor (Fig. 2e) These movements are similar to both the prototypical G_s_-coupled family A receptor, the β_2_-adrenoceptor^16^, and to the LH/CGR^19^ (Supplementary Fig. 6).

A central question is how conformational changes in the TSHR ECD connect to activation of the TSHR 7TM domain. Our structures reveal that the C-terminus of the TSHR ECD hinge region wraps around the hinge helix to connect to the p10 region (Fig. 2d). The hinge region is unresolved in inactive TSHR, demonstrating a disorder-to-order transition upon binding to TSH and supporting an important role for this region in hormone recognition. The hinge helix connects to the rest of the TSHR ECD and the p10 region via two disulfide bonds. Rotation of the TSHR ECD to the up state is coupled to rotation of the hinge helix, which acts like a pulley to displace and lift the p10 region towards the extracellular side. The resulting conformation of the p10 region yields several new contacts with the extracellular loops of TSHR and an inward movement of TM7 that together likely stabilize the active conformation of the 7TM domain.

### Agonist M22 autoantibody mimics TSH to activate TSHR

We next aimed to understand how agonistic autoantibodies activate the TSHR. Previous efforts to characterize thyroid stimulating immunoglobulins (TSI) identified M22, a monoclonal antibody isolated from a patient with Graves’ disease that potently activates the TSHR^36^. A prior X-ray crystal structure of M22 bound to the extracellular domain of the TSHR provided evidence of an epitope overlapping with the predicted binding site for TSH^37^. However, a neutral antagonist antibody with TSH-blocking properties, K1-70, also binds at an almost completely overlapping epitope to M22^38^. The discordant signaling activities of K1-70 and M22 suggest that the interaction between TSIs and the 7TM domain of the TSHR are critical to autoantibody activation, but remain poorly understood^39,38^.

We therefore prepared a complex of active TSHR-miniG_s_ bound to the Fab fragment of the M22 autoantibody and analyzed the structure and dynamics of this complex with cryo-EM. In 2D class averages, we identified two distinct conformations of the TSHR ECD bound to the M22 Fab (Fig. 3a). One class appeared consistent with a TSHR ECD conformation in the up state similar to that observed for TSH. In the other class, the TSH ECD:M22 Fab complex is distinct from either the up or down conformations of the TSHR ECD. Instead, the TSH ECD appears to embed within the side of the detergent micelle in a region that would normally occupy the lipid bilayer. Although this “side” orientation of the TSHR ECD is not physiological and likely results from cryo-EM of detergent-solubilized TSHR, it suggests that the M22-bound TSHR ECD is more conformationally heterogeneous than when bound to TSH. We were unable to determine a high-resolution reconstruction of this “side” complex (Supplementary Fig. 7). However, we successfully reconstructed two high resolution maps for the up state: the TSHR ECD bound to M22 Fab at 3.0 Å and the TSHR 7TM domain bound to Gs at 2.8 Å (Fig. 3b and Supplementary Fig. 7).

**Figure 3.**
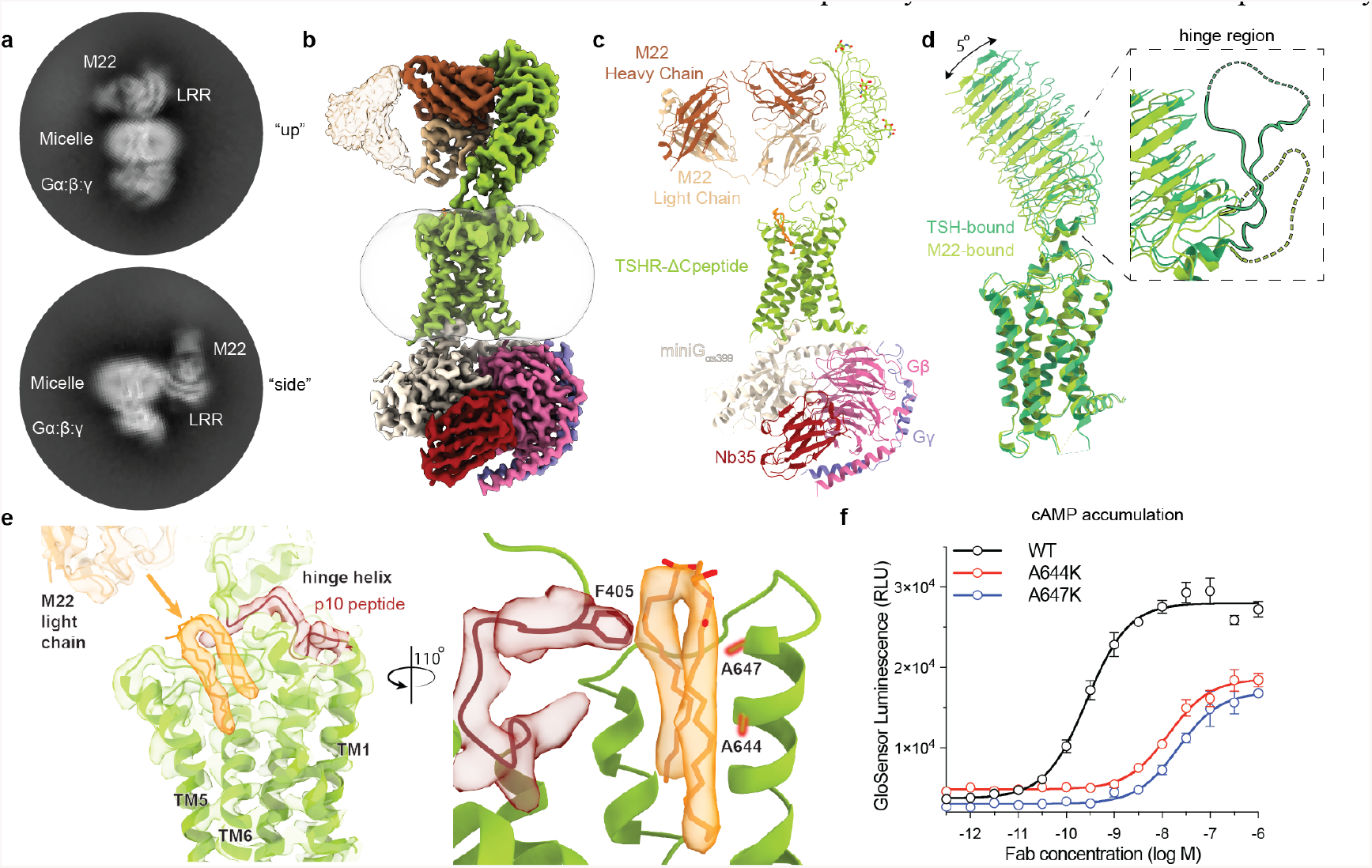
Activation of TSHR by a Graves ’disease autoantibody. **a)** Selected 2D class averages of the “up” and “side” TSHR ECD orientations, highlighting the range of conformational plasticity in ECD orientations upon M22 Fab binding. Cryo-EM map (**b**) and model (**c**) of the M22 Fab-TSHR-G_s_-Nb35 complex. **d)** Alignment of TSHR 7TM domain between TSH and M22-bound models reveals minimal (∼5°) change in the orientation of the ECD but a significant reduction in the resolved density of the M22 hinge region. **e)** DPPC (orange) is modeled into the clear two-tailed density present in the TSHR transmembrane pocket. F405 in the p10 peptide appears to accommodate the hydrophobic tails of DPPC in the active-state conformation of this receptor region. A644 and A647 (red) were individually mutated to K and tested for **f)** changes in M22-mediated cAMP accumulation via the GloSensor assay. Signaling data points represent the global fit of grouped triplicate measurements at each concentration ±SD from 4 independent experiments.

We next compared TSH- and M22-activated TSHR (Fig. 3d). With M22, the ECD up state is in a position that is highly similar to the TSH-activated state, with a small 5 degree rotation when the 7TM domains are aligned. The conformation of the 7TM domain and orientation of G_s_ heterotrimer are also highly similar between M22- and TSH-activated TSHR, with a RMSD of 0.87 Å. A key distinction between TSH- and M22-activated TSHR is that the hinge region is unresolved in M22-activated TSHR, which is consistent with prior studies showing that the hinge region is dispensable for activation of the TSHR^32^ (Fig. 3d, inset). The lack of a direct interaction with the hinge region is likely responsible for the increased conformational flexibility of the TSHR ECD when bound to M22 as compared to TSH.

The higher-resolution reconstruction of M22-bound TSHR enabled us to resolve density for a phospholipid buried within the 7TM domain of the receptor in a region that overlaps with the likely binding side for Org 274179-0 in TSHR and Org43553 in LH/CGR (Fig. 3e and Supplementary Fig. 8). We tentatively assigned this density to arise from dipalmitoylphosphatidylcholine (DPPC). This designation was informed by mass spectrometry experiments to identify potential lipids enriched in our TSHR-G_s_ samples and potential candidates compatible for modeling within the observed cryo-EM density (Supplementary Fig. 8). The well-resolved aliphatic tails of DPPC make extensive transmembrane domain contacts; the headgroup, by contrast, is not resolved. Additionally, DPPC appears to make direct contact with F405 of the p10 region (Fig. 3e). Previous mutagenesis to other hydrophobic residues (T, L) at this site has been shown to increase basal and agonist-mediated cAMP accumulation^40^. Thus, we investigated whether lipid occupancy in this transmembrane pocket is important in activation of the TSHR. Two mutations predicted to inhibit binding of DPPC, A644K and A647K significantly decrease the efficacy and potency of M22 at the TSHR (Fig. 3f). These mutations also have a moderate effect on cell surface expression of the TSHR (Supplementary Fig. 1). This suggests that lipid binding at this site may also play a role in receptor folding and trafficking while also serving as a critical conduit to connect the conformation of the 7TM domain with the p10 region, and thereby to the rotation of the TSHR ECD.

### Membrane bilayer is critical for TSHR activation

Our structures of the TSHR revealed that rotation of the ECD is coupled to 7TM domain activation. A central question, however, is how binding of TSH and activating TSI lead to the up ECD conformation. Based on structures of the LH/CGR, Duan *et. al* proposed that a clash between the CG-β chain and the membrane bilayer drives activation by “pushing” the LH/CGR ECD away from the membrane bilayer^19^. Simultaneously, an additional interaction between the hinge loop and the common GPH-α chain “pulls” the ECD to the up state. While our structures of inactive and active TSHR revealed overall conformation changes similar to LH/CGR, we arrive at a distinct model for glycoprotein hormone action based on our structures of TSH, the M22 antibody agonist, and the CS-17 antibody inverse agonist.

We first modeled how the membrane bilayer may interact with TSH, M22, or CS-17 when bound to the TSHR ECD in either the active up or inactive down conformations (Fig. 4a). Here, we used a computational approach to place the TSHR within a membrane bilayer to most accurately estimate the overall orientation of the receptor^41^ (Fig. 4b). In the inactive state, the down conformation of the TSHR ECD would be unable to bind TSH due to clashes between the glycosylated Asn52 residue in the common GPHα chain. Because typical biantennary N-linked glycans are composed of 5-12 monosaccharide units with an average size of ∼9 Å per monosaccharide^45^, a glycan at Asn52 would be predicted to preclude the down ECD state. Importantly, there is no predicted clash between the protein component of TSH and the membrane bilayer, suggesting that glycosylation at GPHα Asn52 is critically important for TSHR activation. Indeed, prior studies have demonstrated that glycosylation of TSH and CG is necessary for TSHR and LH/CGR activation, respectively. Deglycosylated TSH or CG bind to their respective receptors, but function as competitive antagonists for the native hormone^11,42,43,44^, supporting the critical role that glycosylation plays in glycoprotein hormone activation of their cognate receptors.

**Figure 4.**
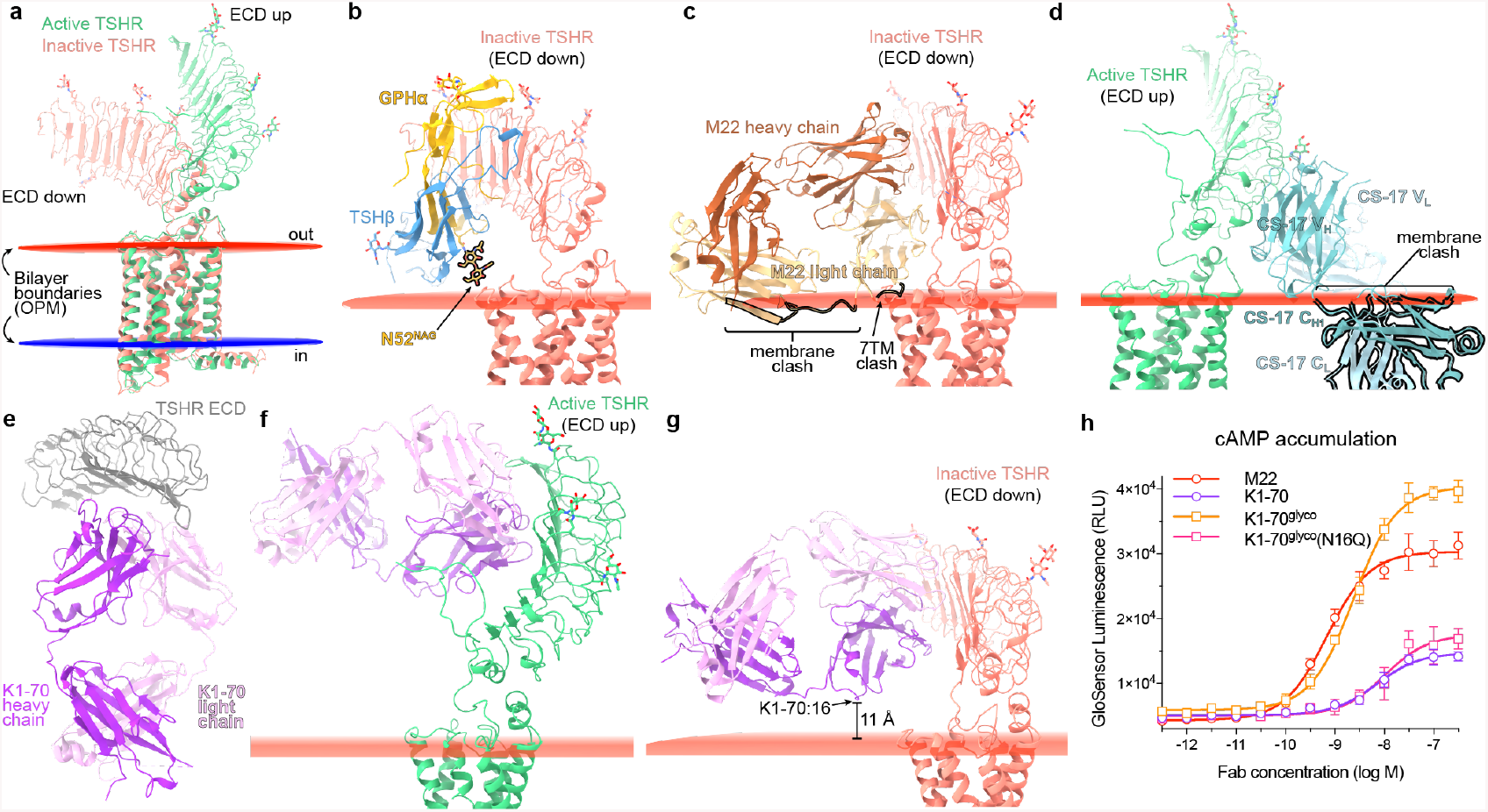
Membrane bilayer interactions are critical for TSHR activation. **a)** Orientation of active and inactive TSHR in a mammalian plasma membrane bilayer as defined by the Orientations of Proteins in Membranes server. Modeling of TSH binding to inactive TSHR with the ECD in the down conformation with either TSH (**b**) or M22 Fab (**c**). The protein components of TSHβ and GPHα are not predicted to clash with the membrane bilayer. Biantennary glycosylation at position GPHα N52 would extend beyond the two N-acetylglucosamine (NAG) units, which would likely clash with the membrane bilayer in inactive TSHR. Loops in the M22 light chain are predicted to clash with both the 7TM domain of inactive TSHR and the membrane bilayer. **d)** Modeling of inverse agonist CS-17 binding to active TSHR. **e)** X-ray crystal structure of K1-70 Fab bound to TSHR ECD (PDB ID: 2XWT). Modeling of K1-70 bound to either active TSHR (**f**) or inactive TSHR (**g**). K1-70 is compatible with binding to both ECD up and ECD down states. Residue 16 in K1-70 heavy chain is 11 Å distant from the membrane. **h)** An engineered version of K1-70 with N-linked glycosylation at residue 16 (K1-70^glyco^) is more potent and efficacious at cAMP accumulation downstream of TSHR than K1-70. Signaling data points represent the global fit of grouped sextuplicate measurements at each concentration ±SD from 3 independent experiments.

We used a similar approach to understand the structural basis for the activity of M22 and CS-17 antibodies. In the inactive down conformation of the TSHR ECD, the M22 light chain would clash with the 7TM domain and the membrane bilayer (Fig. 4c). Conversely, in the active up state of the TSHR ECD, the CS-17 antibody constant domains of both heavy and light chain would clash with the membrane bilayer (Fig. 4d). We surmise that TSH and M22 are both activators because they prevent the conformational transition of the TSHR ECD to the inactive down state and CS-17 is an inverse agonist because it precludes the TSHR ECD active up state.

To directly test this model, we designed a gain-of-function experiment with the K1-70 antibody fragment (Fig. 4e-h). The K1-70 antibody has been characterized as a TSH antagonist; in our hands, it weakly activates the TSHR in a cAMP signaling assay (Fig. 4h). Aligning the previously solved X-ray crystal structure of K1-70 with the inactive state of the TSHR suggests that, unlike M22, K1-70 does not clash with the membrane bilayer (Fig. 4e-g). We engineered a construct, K1-70^glyco^, with a predicted glycosylation motif in a loop that is the closest contact between K1-70 and the membrane bilayer (Fig. 4g). Our prediction was that the engineered glycan would mimic the effect of TSH glycosylation at Asn52 and clash with the membrane bilayer in the inactive down state of TSHR, thereby converting K1-70 into a highly efficacy agonist. We isolated a glycosylated preparation of K1-70^glyco^ by Concanavalin A affinity chromatography and confirmed its glycosylation status by native mass spectrometry (Supplementary Fig. 9). Consistent with our model, K1-70^glyco^ is significantly more efficacious at activating cAMP signaling at TSHR compared to either K1-70 or a non-glycosylated control K1-70^glyco^(N16Q), even surpassing the maximal effect of the potent M22 Fab (Fig. 4h). We conclude that sterically precluding the “down” state of the TSHR ECD is sufficient for receptor activation.

## Discussion

Based on our structural observations and signaling studies, we propose the following model for TSHR activation (Fig. 5). In the basal state, the TSHR ECD is likely structurally dynamic and can transiently exist in the up conformation. These transient excursions lead to basal signaling. TSH selects the ECD up state due to clashes between the GPHα Asn52 glycan and the membrane bilayer if the TSH ECD converts to the down state. The ECD up state activates the 7TM domain via the hinge helix, the p10 peptide, and a phospholipid within the 7TM domain. While interactions between the hinge region and TSH are required for potent hormone binding and signaling, it is not required *per se*, as TSHR is activated by the M22 autoantibody without similar contacts. Our gain-of-function experiments with K1-70 highlight that a single N-linked glycosylation is sufficient to activate TSHR by restricting the TSHR ECD in an up state. Finally, conformational selection of the ECD down state leads to inverse agonist activity. The conformation of the ECD relative to the membrane bilayer is therefore a critical determinant of TSHR activity.

**Fig. 5.**
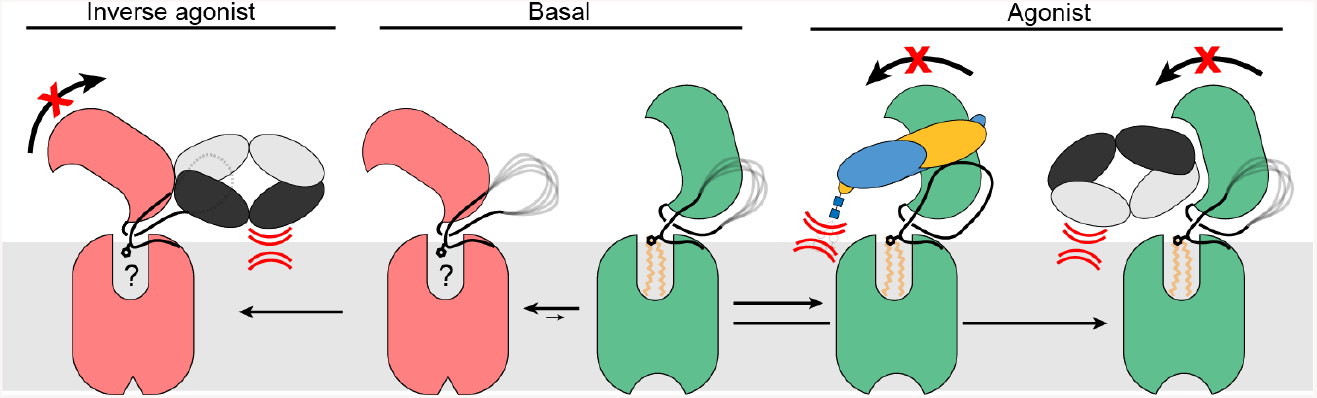
Model for TSHR action. In the basal state, the TSHR ECD can spontaneously transition to the up state, leading to constitutive activity. TSH stabilizes an upright ECD because steric clashes between the GPHα N52 glycan and the membrane bilayer prevent conversion of the ECD to the down state. Agonistic autoantibodies like M22 activate TSHR in a similar manner by preventing the ECD down state. Conversely, inverse agonistic antibodies like CS-17 prevent the ECD from assuming the up state, thereby locking TSHR in an inactive conformation.

Our studies revise current models of glycoprotein hormone receptor activation. We demonstrate that interaction between the hormone and receptor hinge region is not sufficient for receptor activation. Instead, a critical determinant of activity is the membrane interaction between the GPHα Asn52 glycan and the membrane bilayer, which is conserved between all glycoprotein hormones.

Glycosylation at the right position itself is sufficient to activate the TSHR as illustrated by our experiments with K1-70, underscoring the importance of this post-translational modification in glycoprotein hormone action. Our studies with agonistic and inverse agonistic antibodies demonstrate that conformational selection of the ECD is sufficient to activate or inactivate the receptor. Indeed the lower constitutive activity of both LH/CGR and FSHR^46,47^ is likely explained by decreased ECD conformational dynamics in these receptors relative to TSHR. Finally, our structure with M22 reveals a phospholipid embedded within the transmembrane domain of TSHR in a region that is overlapping with existing activating and inhibiting molecules for the FSHR and LH/CGR. We therefore predict that a similar phospholipid may be a common feature that underlies glycoprotein hormone receptor activation.

The structural basis of how autoantibodies mimic the TSH hormone to activate the TSHR provide insight into the molecular pathophysiology of Graves’ disease, and more broadly, highlight the unique way in which breaking of immune tolerance to self-antigens leads to aberrant GPCR signaling. For Graves’ disease in particular, our structure of the TSHR bound to M22 suggests that the progression of thyroid autoimmune diseases is largely dependent on the geometric binding behavior of the TSHR ECD interacting antibodies relative to the membrane bilayer. This provides some context for the presence of both TSH blocking and agonistic autoantibodies in patients with autoimmune thyroid disease - although these antibodies likely share similar epitopes on the TSHR ECD, it is their orientation relative to the membrane bilayer that determines their efficacy. More broadly, our studies provide a first perspective on how autoantibodies pathologically activate GPCRs, of which there are now dozens of examples in the literature^48^. While the specific mechanisms for such pathological antibodies acting at various GPCRs are likely unique, a shared feature is likely recognition of GPCR extracellular domains and conformational selection to mimic hormone function.

Our observations for the TSHR may also have broader implications for GPCR activation by other receptors containing large leucine-rich repeat extracellular domains such as LGR4/5/6 and the relaxin-family receptors. As the LRR domains in these receptors present static interfaces for ligand binding, we suspect that the rigid-body motions and membrane-dependent interactions described for the agonist-bound TSHR ECD are shared amongst these receptors in the transition to activated conformations. Together, our work establishes a key structural role for hormone glycosylation in receptor activation, introduces additional lipid-dependent factors for consideration in the development of small-molecules targeting TSHR, and illuminates the mystery of how both stimulatory and TSH-blocking autoantibodies can exist within a single patient with autoimmune thyroid diseases. Our structural insights set the foundation for discovery of new therapies targeting the TSHR for diseases of thyroid homeostasis.

## Methods

### Expression and purification of TSHR for active state structures

The human *TSHR* gene with an N-terminal influenza hemagglutinin signal sequence and FLAG (DYKDDDK) epitope tag was cloned into a custom pCDNA3.1 vector containing a tetracycline inducible cassette. To improve expression of intact receptor for structural studies, the sequence for the 50-residue hinge-region “C-peptide” (Ala317-Phe366) was removed via site directed mutagenesis (New England Biolabs). The construct further included the miniG_s399_ protein^15^, which was fused to the C-terminus of TSHR with a human rhinovirus 3C protease cleavage sequence flanked by three-residue Gly/Ser linkers. This construct (TSHR-ΔCpep-miniG_s399_) was transfected into inducible Expi293F cells (Thermo Fisher) using the ExpiFectamine transfection reagent per manufacturer instructions. After 24 hours, protein expression was induced with 2 µg/mL doxycycline hyclate, and the culture was placed in a 30°C incubator for 36 hours prior to harvesting by centrifugation. Pelleted cells were washed with 50 mL of phosphate buffered saline, pH 7.5 prior to storage at -80°C. For receptor purification, frozen cells were hypotonically lysed in 50 mM HEPES, pH 7.5, 1 mM EDTA, 160 µg/mL Benzamidine, 2 µg/mL Leupeptin for 10 minutes at 25°C. The membrane fraction was collected by centrifugation, and the fusion protein was extracted with 50 mM HEPES, pH 7.5, 300 mM NaCl, 1% (w/v) glyco-diosgenin (GDN, Anatrace), 0.1% (w/v) cholesteryl hemisuccinate (CHS, Steraloids), 2 mM MgCl_2_, 2 mM CaCl_2_, 160 µg/mL Benzamidine, 2 µg/mL Leupeptin with dounce homogenization and incubation with stirring for one hour at 4°C. The soluble fraction was separated from the insoluble fraction by centrifugation and was incubated in batch for 1 hour at 4°C with homemade M1-FLAG-antibody conjugated sepharose beads. Sepharose resin was then washed extensively with 50 mM HEPES, pH 7.5, 150 mM NaCl, 0.1% (w/v) GDN, 0.01% (w/v) CHS, 2 mM MgCl_2_, 2 mM CaCl_2_ and then with 50 mM HEPES, pH 7.5, 150 mM NaCl, 0.0075% (w/v) GDN, 0.00075% (w/v) CHS, 2 mM MgCl_2_, 2 mM CaCl_2_ prior to elution with 50 mM HEPES, pH 7.5, 150 mM NaCl, 0.0075% (w/v) GDN, 0.00075% (w/v) CHS, 5 mM EDTA, 0.2 mg/mL FLAG peptide. Post-elution, TSHR-ΔCpep-miniG_s399_ fusion protein was concentrated in a 100 kDa MWCO Amicon spin concentrator, and injected onto a Superdex200 Increase 10/300GL (Cytiva) gel filtration column equilibrated in 50 mM HEPES, pH 7.5, 150 mM NaCl, 0.005% (w/v) GDN, and 0.0005% CHS to isolate monodisperse material for further complexing with agonist, Gβ_1_γ_2_ heterodimer, and Nb35^16^.

### Expression and purification of TSHR for inactive state structures

A pCDNA3.1-containing TSHR-ΔCpep construct was generated via site directed mutagenesis. Expi293 cell transfection, doxycycline induction, and expression were performed as described for TSHR-ΔCpep-miniG_s399_ with the additional supplement of dimethylsulfoxide solubilized 1 µM Org 274179-0^33^ post-induction. Purification of TSHR-ΔCpep was performed as described for TSHR-ΔCpep-miniG_s399_ with all buffers supplemented with 1 µM Org 274179-0.

### Expression and purification of Gβ_1_γ_2_

Human Gβ_1_γ_2_ heterodimer was expressed in *Trichoplusia ni* (Hi5) insect cells using a single baculovirus generated in *Spodoptera frugiperda* (Sf9) insect cells. A bicistronic pVLDual construct contained the Gβ_1_ subunit with a N-terminal 6x His-tag, and an untagged human Gγ_2_ subunit. For expression, Hi5 insect cells were transduced with baculovirus at a density of ∼3.0 × 10^6^ cells/mL, grown at 27 °C shaking at 130 rpm. 48 hours post-transduction, cells were harvested and washed in a hypotonic buffer containing 20 mM HEPES, pH 8.0, 5 mM β-mercaptoethanol (β-ME), and protease inhibitors (20 µg/mL leupeptin, 160 µg/mL benzamidine). The membrane fraction was then separated by centrifugation and solubilized with 20 mM HEPES pH 8.0, 100 mM sodium chloride, 1.0% sodium cholate, 0.05% dodecylmaltoside (Anatrace), and 5 mM β-mercaptoethanol (β-ME). Solubilized Gβ_1_γ_2_ heterodimer was then incubated with HisPur™ Ni-NTA resin (Thermo Scientific) in batch. Bound Gβ_1_γ_2_ heterodimer was washed extensively and detergent was slowly exchanged to 0.1% (w/v) lauryl maltose neopentyl glycol (L-MNG, Anatrace) and 0.01% CHS before elution with 20 mM HEPES pH 7.5, 100 mM NaCl, 0.1% L-MNG, 0.01% CHS, 270 mM imidazole, 1 mM dithiothreitol (DTT), and protease inhibitors. Eluted Gβ_1_γ_2_ heterodimer was pooled and rhinovirus 3C protease was added to cleave the N-terminal 6x His-tag during overnight dialysis in 20 mM HEPES pH 7.5, 100 mM NaCl, 0.02% L-MNG, 0.002% CHS, 1 mM DTT, and 10 mM imidazole. To remove uncleaved Gβ_1_γ_2_, dialyzed material was incubated with HisPur™ Ni-NTA resin in batch. The unbound fraction was then incubated for 1 hour at 4 °C with lambda phosphatase (New England Biolabs), calf intestinal phosphatase (New England Biolabs), and antarctic phosphatase (New England Biolabs) for dephosphorylation. Final anion exchange chromatography was performed using a MonoQ 4.6/100 PE (Cytiva) column to purify only geranylgeranylated heterodimer. The resulting protein was pooled and dialyzed overnight in 20 mM HEPES pH 7.5, 100 mM NaCl, 0.02% L-MNG, and 100 µM TCEP, and concentrated with a 3 kDa centrifugal concentrator to a final concentration of 162 µM. Glycerol was added to a final concentration of 20%, and the protein was flash frozen in liquid nitrogen and stored at -80 °C until further use.

### Expression and purification of Nb35

A pET-26b vector containing the Nb35 sequence with a carboxy-terminal Protein C affinity tag (EDQVDPRLIDGK) was transformed into BL21 Rosetta *Escherichia coli* cells and inoculated into 8 L of Terrific Broth supplemented with 0.1% glucose, 2 mM MgCl_2_, and 50 µg/mL kanamycin. Cells were induced with 400 µM IPTG at OD_600_ = 0.6 and allowed to express at 20 °C for 21 hours. Harvested cells were incubated SET Buffer (200 mM Tris pH 8.0, 500 mM sucrose, 0.5 mM EDTA) in the presence of protease inhibitors (20 µg/mL leupeptin, 160 μg/mL benzamidine) and benzonase. To initiate hypotonic lysis, 2 volumes of deionized water were added to the cell mixture after 30 minutes of SET buffer mixing. Following lysis, NaCl was added to 150 mM, CaCl_2_ was added to 2 mM, and MgCl_2_ was added to 2mM and lysate was spun down to remove the insoluble fraction. Supernatant was incubated with homemade anti-Protein C antibody-coupled sepharose. Nb35 was eluted with 20 mM HEPES pH 7.5, 100 mM NaCl, and 2 mM CaCl_2_, 0.2 mg/mL Protein C peptide, and 5 mM EDTA pH 8.0, concentrated in a 10 kDa MWCO Amicon filter and injected over a Superdex S75 Increase 10/300 GL column (Cytiva) size-exclusion chromatography column equilibrated in 20 mM HEPES pH 7.5, 100 mM NaCl. Monodisperse Nb35 fractions were pooled, concentrated, and supplemented with 20% glycerol prior to flash freezing in liquid nitrogen for storage at -80°C until further use.

### Expression and purification of M22 agonist Fab

The M22 heavy and light chains^50^ were cloned into pFastBac with a GP67 signal peptide and N-terminal FLAG epitope and a P2A self-cleaving peptide sequence between the heavy and light chains. The construct also included a C-terminal 8x Histidine epitope tag on the heavy chain. Baculvorius was generated using the Bac-to-bac method (Thermo Fisher), and used to transduce *Trichoplusia ni* (Hi5) insect cells (Expression Systems) at a density of 2.0 × 10^6^ cells/mL. Cells were cultured with shaking at 120 RPM at 25°C for 60 hours. M22 Fab was purified from the cell supernatant by first adjusting pH to 8.0 with 1 M Tris prior to a 1 hour incubation with 5 mM CaCl_2_, 5 mM MgCl_2_, and 1 mM NiCl_2_ to precipitate chelators. After separating the insoluble fraction, the 8x-His tagged M22 Fab was captured on HisPur™ Ni-NTA resin, washed in a buffer comprised of 50 mM HEPES, pH 7.5, 500 mM NaCl, 20 mM imidazole, pH 7.5 and eluted with the same buffer supplemented with 500 mM imidazole. The eluate was further purified by size exclusion chromatography over a Superdex200 Increase 10/300GL gel filtration column equilibrated in 50 mM HEPES, 100 mM NaCl, pH 7.5. Monodisperse M22 Fab fractions were pooled, concentrated, and supplemented with 20% glycerol prior to flash freezing in liquid nitrogen for storage at -80°C until further use.

### Expression and purification of WT K1-70, K1-70^glyco^ and K1-70^glyco^(N16Q) Fabs

Heavy and light chain sequences for K1-70 IgG^38^ were cloned into a pCDNA3.1 vector containing human IgG1 constant regions. Resulting constructs were transfected into Expi293F cells using a 2:1 mass ratio of heavy to light chain using the Expifectamine transfection kit, per the manufacturer’s instructions. After 5 days, cultures were harvested and the supernatant purified over a 1 mL MabSelect SuRe HiTrap column (Cytiva) equilibrated in 100 mM sodium phosphate, 150 mM sodium chloride, pH 7.2. Monoclonal antibodies were eluted with 100 mM Glycine, pH 3.0, neutralized with 1.5 M Tris pH 8.0, and dialyzed into 20mM sodium phosphate, 10 mM EDTA, pH 7.0 for generation of Fab fragments. Immobilized papain agarose was equilibrated in the IgG dialysis buffer with freshly added 20 mM Cysteine-HCl, pH adjusted to 7.0. IgG was concentrated to ∼40 mg/mL with a 100 kDa MWCO Amicon filter, diluted 1:1 with the IgG dialysis buffer plus Cysteine-HCl, and added to the equilibrated papain agarose. The IgG-papain suspension was placed on a shaker at 37C for overnight digestion. Post-digest, -Fc fragments were removed via 1 hour, room temperature batch incubation with phosphate buffered saline-equilibrated Protein A agarose (Pierce). Fab fragments were concentrated in a 10 kDa MWCO Amicon filter and injected onto a Superdex200 Increase 10/300GL gel filtration column equilibrated in 50 mM HEPES, 100 mM NaCl, pH 7.5. Monodisperse Fab fractions were pooled, and digestion/-Fc removal was confirmed by SDS-PAGE.

To produce a glycosylated version of K1-70 (K1-70^glyco^), a glycosylation motif was introduced at position 16 in the heavy chain in a loop region that connects the A and B strands of the VH IgG domain. The sequence “KKPGQS” was replaced with “KKPGNGS” to generate an N-linked N-x-S/T glycosylation motif. As a control for glycosylation, we also generated a version of K1-70^glyco^ with a N16Q mutation K1-70^glyco^(N16Q). The resulting constructs were then purified as full length IgG as described above for K1-70. Despite introduction of the glycosylation motif in K1-70^glyco^, the extent of glycosylation was not complete. To enrich for the glycosylated fraction of K1-70^glyco^, monodisperse K1-70^glyco^ Fab was brought to 1 mM Mn^2+^ and Ca^2+^ and loaded over a Concanavalin A-conjugated Sepharose 4B packed HiTrap column (Cytiva) pre-equilibrated in 20 mM HEPES, 500 mM NaCl, 1 mM Mn^2+^, 1 mM Ca^2+^, pH 7.4. The glycosylated fraction K1-70^glyco^ was eluted with 20 mM HEPES, 500 mM NaCl, 500 mM methyl alpha-D-mannopyranoside, pH 7.4, fractions were concentrated to 100 µM and flash frozen in liquid nitrogen for storage at -80°C until further use.

### Expression and purification of CS-17 Fab

Heavy and light chain sequences of CS-17 were determined from sequencing of CS-17 murine hybridoma cell line PTA-8174 (ATCC) by Genscript. The resulting sequences were cloned into a pCDNA3.4-containing murine IgG2a construct. Heavy and light chain constructs were transfected, expressed and purified as previously described for K1-70 IgG. CS-17 Fab generation also followed identical steps to K1-70 Fab generation.

### Preparation of inactive- and active-state TSHR complexes

To prepare the TSH and M22-activated TSHR complexes, purified TSHR-ΔCpep-miniG_s399_ was incubated with a 2-fold molar excess of purified Gβ_1_γ_2_, Nb35, and either M22 Fab or native human TSH (National Hormone and Pituitary Program) and incubated overnight at 4 °C. After incubation, the complexed material was purified with anti-Protein C antibody Sepharose resin to purify Nb35-bound complex. Protein C Sepharose was washed with 20 column volumes of 50 mM HEPES, 150 mM NaCl, 0.0075% GDN (w/v), 2 mM CaCl_2_ prior to elution with 50 mM HEPES, 150 mM NaCl, 0.005% GDN (w/v), 5 mM EDTA, 0.2 mg/mL Protein C peptide. The eluted fractions were concentrated with a 100 kDa MWCO Amicon filter, and injected onto a Superdex200 Increase 10/300GL gel filtration column equilibrated in 50 mM HEPES, 150 mM NaCl, 0.005% GDN (w/v), pH 7.5. Monodisperse fractions were concentrated with a 100 kDa MWCO Amicon filter immediately prior to cryo-EM grid preparation.

For formation of the inactive state CS-17-bound TSHR-ΔCpep complex, the CS-17 Fab was incubated at a 2x molar excess with SEC-purified TSHR-ΔCpep overnight at 4 °C, then concentrated in a 100 kDa MWCO Amicon filter prior to cryo-EM grid preparation prior to size exclusion chromatography over a Superdex200 Increase 10/300GL gel filtration column equilibrated in 50 mM HEPES, 150 mM NaCl, 0.005% GDN (w/v), pH 7.5 and 1 µM Org 274179-0. Monodisperse fractions of the resulting complex were concentrated with a 50 kDa MWCO concentrator immediately prior to cryo-EM grid preparation.

### Cryo-EM vitrification, data collection, and processing

#### TSH-bound TSHR-G_s_ complex

The TSH-bound TSHR-G_s_ complex was concentrated to 23 µM and 3 µL was applied onto a glow discharged 300 mesh 1.2/1.3 gold grid covered in a holey gold film (UltrAufoil). After a 30 second hold, excess sample was removed with a blotting time of 3 seconds with a blotting force of 0 prior to plunge freezing into liquid ethane using a Vitrobot Mark IV (Thermo Fisher). 15,346 super-resolution videos were recorded with a K3 detector (Gatan) on a Titan Krios (Thermo Fisher) microscope operated at 300 keV with a BioQuantum post-column energy filter set to a zero-loss energy selection slit width set of 10 eV. 66-frame videos were recorded for 2 s at a nominal magnification of 130,000x (physical pixel size of 0.664 Å /pixel) and a defocus range of -0.8 to - 2.2 µm for a total dose of 77 e-/A^2^.

Super-resolution videos of the TSH-TSHR-G_s_ complex were motion corrected, binned to physical pixel size, and dose fractionated on-the-fly during data collection using UCSF MotionCor2^51^. Corrected micrographs were imported into cryoSPARC^52^ for CTF estimation via the “Patch CTF Estimation” job type. Templates for particle picking were generated from projections of the same TSHR complex reconstructed from a previous 200 keV imaging session. Particle picking templates were low-pass filtered to 20 Å and used to pick 9,151,778 particles. After picking, estimated CTF fit resolution >5 Å and relative ice thickness outlier measurements were used to remove low-quality micrographs before further processing. 6,185,950 curated particles were extracted in a 512-pixel box and Fourier cropped to 128 pixels before undergoing a round of 3D classification with alignment utilizing one 2 0Å low-pass filtered reconstruction and three “random” reconstructions generated from a prematurely truncated *Ab initio* reconstruction job. 2,474,380 particles classified into the “TSHR” class were extracted in a 512-pixel box and Fourier cropped to 256 pixels for two additional rounds of 3D classification with alignment, utilizing the same class distributions as previously described. From these 2 rounds of 3D classification, and because the TSHR ECD remained poorly resolved, 734,891 particles belonging to the “TSHR” class were extracted into a 512-pixel box for a new 3D classification workflow. First, these particles were subject to one round of “non-uniform” refinement^53^ in cryoSPARC and then classified utilizing two different masking schemes (TSHR 7TM domain or TSHR ECD:TSH complex) using alignment-free 3D classification in RELION^54^. Particles in qualitatively “good” looking classes were re-imported into cryoSPARC for focused refinements. Particles from the TSHR 7TM domain-masked 3D classification were subject to a round of focused refinement with a mask encompassing the TSHR 7TM domain and G protein/Nb35 complex. Particles from the TSHR ECD:TSH complex masked-3D classification were subject to two rounds of focused refinement using the same mask as in RELION, decreasing mask pixel dilation between rounds. For each focused refinement, pose/shift gaussian priors (7° standard deviation of prior (rotation), 4Å standard deviation of prior (shifts)) were used to limit large deviations from the initially determined poses. Directional FSC curves for each final reconstruction were calculated, and each map was loaded into ChimeraX^55^ for generation of a composite map via the *vop maximum* command.

#### M22-bound TSHR-G_s_ complex

The M22-bound TSHR-G_s_ complex was concentrated to 13.8 µM and 3 µL was applied onto a glow discharged 300 mesh 1.2/1.3 gold grid covered in a holey gold film and plunge frozen into liquid ethane utilizing identical blotting procedures as with the TSH complex. 25,030 super-resolution videos were recorded with a K3 detector on a 300 kV FEI Titan Krios microscope located at the HHMI Janelia Research Campus equipped with a spherical aberration corrector and a post-column BioQuantum energy filter set to a zero-loss energy selection slit width set of 20 eV. 60-frame videos were recorded across a defocus range of -0.8 to - 2.0 µm at a nominal magnification of 81,000x (physical pixel size of 0.844 Å /pixel) for 4.4s, resulting in a total exposure dose of 60 e-/A^2^.

25,030 super-resolution videos were motion corrected, binned to physical pixel size and dose weighted, post acquisition, using UCSF MotionCor2. Contrast transfer function estimation was performed with the cryoSPARC “Patch CTF Estimation” job prior to automated particle picking with a gaussian template. After curating for high quality micrographs as previously described, 9,078,725 particles were extracted in a 512-pixel box and Fourier cropped to 128 pixels. Due to the size of the dataset, image processing was performed in three cohorts. In the first cohort, 2,978,754 particles were subjected to two rounds of 2D classification followed by *Ab initio* reconstruction of selected particles. Distinct conformations of the TSHR ECD in “up” and “side” states were apparent from 2D classification. Particles were selected for one round of 3D classification without alignment irrespective of their ECD conformation, then split into a second separate round 3D classification in “up” or “side” classes only. Particles from the best looking and highest resolution “side” class were extracted in an unbinned 512 pixel box and subject to non-uniform refinement. The resulting reconstruction was of low quality, with uninterpretable 7TM domains that contoured at thresholds similar to the detergent micelle. As a result, particle images belonging to “side” class were not further processed. Particles classified into a high-quality ECD “up” class were extracted into a 512 pixel box and subject to non-uniform refinement. The remaining 16,159 micrographs were processed in a similar workflow of multiple rounds of 2D classification followed by 3D classification with alignment. Unbinned particles from each cohort in qualitatively good ECD “up” classes were subject to independent non-uniform refinement jobs. The three refined sets of particles were then combined and subject to unmasked, alignment-free 3D classification in RELION. All classes exhibited high quality 7TM domain features.

Thus, particles in classes containing distinct ECD:M22 VH/VL density were selected, combined, and re-imported into cryoSPARC for 1 round of focused refinement using masked regions separately encompassing the TSHR ECD:M22 domain or the TSHR transmembrane:G protein complex. Directional FSC curves for each focused refinement reconstruction were calculated, and each map was loaded into ChimeraX for generation of a composite map via the *vop maximum* command.

#### Org 274179-0-bound TSHR complex

The Org 274179-0-bound TSHR-ΔCpep complex was concentrated to 55 µM and 3 µL was plunge frozen on holey gold grids as previously for other TSHR samples described after a 30 second hold and a 3 second blotting time at a blotting force of 0. Cryo-EM data were collected on a 300 keV Titan Krios microscope equipped with a K3 detector and a BioQuantum post-column energy filter with a slit width of 20 eV. 10,003 super-resolution movies were recorded at a nominal magnification of 105,000x (physical pixel size of 0.86 Å /pixel) for 1.35 s each and fractionated across 50 frames. The defocus range for this collection was -0.8 to - 2.4 µm.

10,003 TSHR-ΔCpeptide-Org 274179-0 videos were motion corrected, binned to physical pixel size and dose weighted, post acquisition, using UCSF MotionCor2. CTF estimation and reference-free “blob” particle autopicking were performed in cryoSPARC. Micrographs were curated following similar thresholds as previously described prior to extraction of 5,922,556 particles in a 288-pixel box that was subsequently Fourier cropped to 72 pixels. To generate a suitable reference for 3D classification, 2 rounds of 2D classification were performed. From 2D classification, the particles belonging to classes that exhibited “micelle”-like density were then subjected for *Ab initio* 3D reconstruction. Then, 3D classification with alignment was performed on the entire set of initially picked particles using one class suggestive of a micelle:ECD reconstruction and three classes generated from prematurely truncated *Ab initio* jobs, as previously described. Particles classified into the micelle:ECD class were subject to two further rounds of the same classification workflow prior to being extracted without Fourier cropping. Finally, the 357,869 remaining particles were subject to non-uniform refinement. The reconstruction quality of the TSHR-Org 274179-0 complex was poor, and was not improved upon further rounds of unbinned 3D classification with, or without, alignment. 2D classification of the particles in the initial non-uniform refinement job revealed clear density for the ECD with more diffuse alignment on the 7TM domain.

#### CS-17-bound TSHR complex

3µL of purified CS-17-bound TSHR-ΔCpep:Org 274179-0 complex was concentrated to 40.8 µM and similarly plunge frozen into liquid ethane after a 30 second hold, a blotting time of 2 seconds, and a blotting force of 0. The TSHR-ΔCpeptide:Org 274179-0 complex at 55 µM was plunge frozen on holey gold grids as previously described after a 30 second hold and a 3 second blotting time at a blotting force of 0.

9,244 super-resolution videos were recorded with identical collection parameters on the same Titan Krios microscope as described for the TSHR:Org 274179-0 complex.

Next, super-resolution videos of the TSHR-ΔCpeptide:CS-17 Fab fragment complex were motion corrected, dose-weighted, and binned on-the-fly with MotionCor2 during data collection, as previously described. CTF estimation, “blob” particle autopicking, and Fourier cropped particle extraction were all performed in cryoSPARC, after curating for high quality micrographs. 2,742,080 picked particles were subject to 4 rounds of 2D classification followed by *Ab initio* reconstruction to generate reference maps for 3D classification. Then, all picked particles were subject to 4 rounds of 3D classification with alignment, two rounds at a 4x-Fourier cropped box size of 112 pixels, and two rounds at a 2x-Fourier cropped box size. 114,707 particles were then subject to 3D classification with alignment using four identical input reference volumes. The shifts and poses of the 41,054 particles belonging to the highest resolution reconstruction from this final 3D classification were then refined using the non-uniform refinement job type. In an attempt to improve reconstruction quality in the TSHR-ΔCpep:CS-17 Fab binding interface, 2 rounds of focused refinement were performed. The first focused refinement utilized a mask encompassing only the ECD and CS-17 variable domains and was followed by a refinement with the entire TSHR-ΔCpep:CS-17 complex masked. Both focused refinements used the same rotation/shift prior restrictions as in the TSHR-G_s_ processing workflow. However, in comparison to the initial non-uniform refinement reconstruction, the focused refinements yielded lower quality 7TM domain reconstructions. As a result, the pre-focused refinement reconstruction half maps were used for directional FSC calculation and subsequent atomic model building.

### Model Building and refinement

We first modeled the M22-bound TSHR-Gs complex in a composite cryo-EM map (Supplementary Fig. 7). For TSHR, we started with an AlphaFold^56^ model of full-length human TSHR, which had high structural agreement with a previously determined X-ray crystal structure of the M22 Fab bound to the TSHR ECD (PDB ID: 3G04^37^). After truncating unresolved regions, the 7TM domain in this model was fit into the composite cryo-EM map of M22-bound TSHR-Gs with ChimeraX^55^. This template model was rebuilt in Coot^57^ for rigid body fitting of the extracellular domain. For the M22 Fab, we used the PDB ID: 3G04 as a starting structure. For the G protein (miniG_s_, Gβ, and Gγ) we used PDB ID: 7LJC^58^ as a starting template. Finally, for Nb35, we used PDB ID: 3SN6^16^ as a starting template. For DPPC, restraints were generated using the Prodrg server^59^ and the aliphatic tails of the lipid were manually docked into cryo-EM density using Coot. Each of these components was individually fit into cryo-EM density with ChimeraX. We subsequently iteratively refined the model with manual refinement in Coot and ISOLDE^60^ and real space refinement in Phenix^61^.

To model TSH-bound TSHR, we started with the M22-activated structure described above and fit it into a composite map of the TSH-TSHR-G_s_ complex using ChimeraX (Supplementary Fig. 2). The fit of the TSHR ECD was further optimized by rigid body fitting in Coot and Phenix. A key distinction between the M22- and TSH-activated structures of TSHR is in the hinge region. For TSHR, we were able to resolve residues 291-302 and 387-396 in the hinge domain, which were built as a polyalanine chain. To model TSH, we generated a model of the GPHα-TSHβ using AlphaFold, which was fit into the cryo-EM density map in ChimeraX. The resulting model was iteratively refined with manual changes in Coot and ISOLDE and real space refinement in Phenix.

For the CS-17/Org 274179-0 bound TSHR complex, we started with AlphaFold predictions for the TSHR and for the CS-17 Fab which were fit into the cryo-EM map with ChimeraX.The TSHR ECD was fit into the cryo-EM density by rigid body refinement using Coot. Similarly, the constant regions of the CS-17 Fab were fit into the cryo-EM density by rigid body refinement in Coot. The resulting model was refined in ISOLDE with manual changes in both ISOLDE and Coot followed by real space refinement in Phenix.

All maps and models were validated using MolProbity^62^.

### Identification of lipid in M22-bound TSHR-G_s_

Lipids were extracted using a modified version of the Bligh-Dyer method^63^. Briefly, samples or PC(16:0/16:0) (DPPC) for the standard curve were manually shaken for 30s in a glass vial (VWR) with 1 mL PBS, 1 mL methanol and 2 mL chloroform containing the internal standard Cer(d18:1-d7/18:1). The resulting mixtures were vortexed for 15s and centrifuged at 2400 x g for 6 min to induce phase separation. The organic (bottom) layer was retrieved using a glass pipette, dried under a gentle stream of nitrogen, and reconstituted in 2:1 chloroform:methanol for LC/MS analysis. Targeted lipidomic analysis was performed on a Dionex Ultimate 3000 LC system (Thermo) coupled to a TSQ Quantiva mass spectrometer (Thermo). Data was acquired in positive ionization mode. Solvent A consisted of 95:5 water:methanol, Solvent B was 70:25:5 isopropanol:methanol:water. Solvents A and B contained 5 mM ammonium formate with 0.1% formic acid. A XBridge (Waters) C8 column (5 μm, 4.6 mm × 50 mm) was used. The gradient was held at 0% B between 0 and 5 min, raised to 20% B at 5.1 min, increased linearly from 20% to 100% B between 5.1 and 35 min, held at 100% B between 35 and 40 min, returned to 0% B at 40.1 min, and held at 0% B until 50 min. Flow rate was 0.1 mL/min from 0 to 5 min, 0.3 mL/min form 5.1 to 50 min. MS analyses were performed using electrospray ionization in positive ion mode, with spay voltages of 3.5 kV, ion transfer tube temperature of 325 C, and vaporizer temperature of 200 C. Sheath, auxiliary, and sweep gases were 40, 10 and 1, respectively. Internal standard Cer(d18:1-d7/18:1) was detected using the following transitions: 571.6>271.2, 571.6>289.3, 571.6>259.3. DPPC was detected using the following transition: 734.6>184.1, 734.6>125.1, 734.6>86.1. Other PCs were detected using the specific parent ion m/z and the same diagnostic fragments as DPPC. Chromatography and peak integration of lipid targets were verified with Skyline^64^. Peak areas were used in data reporting, data was normalized using internal standards.

### Native mass spectrometry (nMS) of K1-70 fab fragments

Purified Fab fragments were prepared for native mass spectrometry (nMS) by buffer exchange into 200 mM ammonium acetate with a Zeba spin desalting column. Samples, normalized to 5 µM, were injected directly in a Q Exactive Extended Mass Range mass spectrometer using nanoES electrospray capillaries. The instrument parameters were: capillary voltage 1.1 kV; S-lens RF 100%; quadrupole selection 300-10,000 m/z, collisional activation in the HCD cell 150 °C, trapping gas pressure setting 8; temperature 250 °C, instrument resolution 35,000. Data was analyzed with UniDec 4.2.2^65^.

### G_s_ signaling assays

To measure ligand-dependent activation of G_s_ by TSHR we measured cAMP accumulation. For each TSHR construct (wild-type, ΔC-peptide, Y385F, Y385A, A644K, A647K), a 20 mL suspension culture of Expi293 cells was co-transfected with a pCDNA3.1 plasmid expressing TSHR and a luciferase-based cAMP biosensor (pGlosensor-22F; Promega). A total of 20 µg of DNA was transfected using Expifectamine per manufacturer instructions in a 3:1 ratio of receptor construct to cAMP biosensor. Cells were harvested 24 h post-transfection, resuspended in Expi293 expression media supplemented with 10% DMSO, and gradually frozen to -80 °C in a Mr. Frosty Freezing container for future use. To perform the assay, frozen Expi293 cells were rapidly thawed in a 37°C water bath and resuspended in fresh Expi293 expression medium. Cells were diluted to a final concentration of 500,000 cells/mL in Expi293 expression medium plus 2% (v/v) Glosensor reagent (Promega) and incubated for 75 minutes at room temperature with gentle rotation. Expi293 cells were then plated into a white 384-well plate (Greiner) to a final density of 10,000 cells per well. Separately, a 5x ligand stock plate (13-point half log dilution series, 3-wells per condition) was made in Hank’s balanced salt solution + 0.1% (w/v) bovine serum albumin. The ligand plate was stamped into new 384 well plates to achieve a final 1X concentration upon addition of Glosensor-reagent incubated, co-transfected TSHR:cAMP biosensor cells. Immediately after cell addition, luminescence was measured in 0.1s intervals for 10 minutes using a CLARIOstar instrument at an emission wavelength of 580 nm ± 80 nm band-pass (BMG LabTech). The resulting dose response curves from the 5-minute read point were fit to a nonlinear regression three-parameter log(agonist) vs. response fit in GraphPad Prism 9.

## Data Availability

Coordinates for TSH-bound TSHR-G_s_, M22-bound TSHR-G_s_, and CS-17-bound TSHR have been deposited in the PDB under accession codes 7T9I, 7T9N and 7T9M, respectively. Unsharpened EM density maps for TSH-bound TSHR-G_s_ (composite), M22-bound TSHR-G_s_ (composite), and CS-17-bound TSHR have been deposited in the Electron Microscopy Data Bank under accession codes 25758, 25763, and 25762, respectively. Final particle stacks and .star files containing particle shift/pose assignments, half maps for each component of the composite maps (TSH-bound and M22-bound TSHR-G_s_), and half maps for the CS-17-bound TSHR final map have been uploaded to the Electron Microscopy Public Image Archive under the accession code XXXX, XXXX, and XXXX, respectively.

## Acknowledgments

We thank Rui Yan at the HHMI Janelia CryoEM Facility and Dan Toso at Cal-Cryo at QB3-Berkeley for help in microscope operation and data collection, Cole Bracken for mass spectrometry troubleshooting, and Hazel Shan for sample preparation for lipid mass spectrometry. This work was supported by National Institutes of Health (NIH) grants DP5OD023048 (A.M.), 1R35GM140847 (Y.C.), P30CA014195, R01GM102491 (A.S.). Cryo-EM equipment at UCSF is partially supported by NIH grants S10OD020054 and S10OD021741. Some of this work was performed at the Stanford-SLAC Cryo-EM Center (S2C2), which is supported by the National Institutes of Health Common Fund Transformative High-Resolution Cryo-Electron Microscopy program (U24 GM129541). This research was, in part, supported by the National Cancer Institute’s National Cryo-EM Facility at the Frederick National Laboratory for Cancer Research under contract HSSN261200800001E. Some of this work was supported by the Mass Spectrometry Core of the Salk Institute with funding from NIH-NCI CCSG: P30 014195 and the Helmsley Center for Genomic Medicine. The content is solely the responsibility of the authors and does not necessarily represent the official views of the National Institutes of Health. Y.C. is an Investigator of Howard Hughes Medical Institute. A.M. acknowledges support from the Pew Charitable Trusts, the Esther and A. & Joseph Klingenstein Fund and the Searle Scholars Program.

## Author contributions

B.F. cloned, expressed, and biochemically optimized the purification of all TSHR constructs. B.F. expressed and purified Gβ_1_γ_2_, Nb35, M22 Fab, CS-17 IgG, and K1-70 IgGs, and performed enzymatic digestion and further purification to form CS-17 and K1-70 Fab fragments. B.F. performed complexing and identified optimal cryo-EM grid preparation procedures, screened samples, and collected 300 keV datasets. B.F. determined high resolution cryo-EM maps by extensive image processing under the guidance of A.M. and Y.C. B.F. and A.M built and refined models of TSHR complexes. B.F. determined receptor expression levels by flow cytometry and performed and analyzed data from signaling studies. I.S. performed initial biochemical optimization of TSHR complexes with M22 Fab and worked with K.Z. to collect negative stain and cryo-EM data. N.H. prepared samples for, performed, and analyzed native mass spectrometry experiments. N.H. prepared control samples for lipid identification experiments. Y.M. generated TSHR and M22 Fab expression constructs and performed pilot biochemical purification of TSHR complexes. C.B.B. generated expression vectors for WT K1-70 constructs. A.M.P. performed and analyzed data from the lipid identification experiments with guidance from A.S. Figures were generated and the manuscript written by B.F and A.M., with edits from Y.C. and with approval from all authors. The overall project was supervised by Y.C. and A.M.

## References

1. Maenhaut, C. et al. Ontogeny, Anatomy, Metabolism and Physiology of the Thyroid. in Endotext (eds. Feingold, K.R. et al.) (MDText.com, Inc., 2015).

2. Segerson, T. P. et al. Thyroid hormone regulates TRH biosynthesis in the paraventricular nucleus of the rat hypothalamus. Science 238, 78–80 (1987).

3. Fekete, C. & Lechan, R. M. Central regulation of hypothalamic-pituitary-thyroid axis under physiological and pathophysiological conditions. Endocr. Rev. 35, 159–194 (2014).

4. Laurent, E., Mockel, J., Van Sande, J., Graff, I. & Dumont, J. E. Dual activation by thyrotropin of the phospholipase C and cyclic AMP cascades in human thyroid. Mol. Cell. Endocrinol. 52, 273–278 (1987).

5. Taylor, P. N. et al. Global epidemiology of hyperthyroidism and hypothyroidism. Nat. Rev. Endocrinol. 14, 301–316 (2018).

6. Zimmermann, M. B. & Boelaert, K. Iodine deficiency and thyroid disorders. Lancet Diabetes Endocrinol 3, 286–295 (2015).

7. Mincer, D. L. & Jialal, I. Hashimoto Thyroiditis. In StatPearls (StatPearls Publishing, 2021).

8. Lane, L. C., Cheetham, T. D., Perros, P. & Pearce, S. H. S. New Therapeutic Horizons for Graves’ Hyperthyroidism. Endocr. Rev. 41, (2020).

9. Smith, T. J. & Hegedüs, L. Graves’ Disease. N. Engl. J. Med. 375, 1552–1565 (2016).

10. Flack, M. R., Froehlich, J., Bennet, A. P., Anasti, J. & Nisula, B. C. Site-directed mutagenesis defines the individual roles of the glycosylation sites on follicle-stimulating hormone. J. Biol. Chem. 269, 14015–14020 (1994).

11. Matzuk, M. M., Keene, J. L. & Boime, I. Site specificity of the chorionic gonadotropin N-linked oligosaccharides in signal transduction. J. Biol. Chem. 264, 2409–2414 (1989).

12. Grossmann, M., Weintraub, B. D. & Szkudlinski, M. W. Novel insights into the molecular mechanisms of human thyrotropin action: structural, physiological, and therapeutic implications for the glycoprotein hormone family. Endocr. Rev. 18, 476–501 (1997).

13. Chazenbalk, G. D. et al. Evidence that the thyrotropin receptor ectodomain contains not one, but two, cleavage sites. Endocrinology 138, 2893–2899 (1997).

14. Chen, C.-R., Salazar, L. M., McLachlan, S. M. & Rapoport, B. Deleting the Redundant TSH Receptor CPeptide Region Permits Generation of the Conformationally Intact Extracellular Domain by Insect Cells. Endocrinology 156, 2732–2738 (2015).

15. Nehmé, R. et al. Mini-G proteins: Novel tools for studying GPCRs in their active conformation. PLoS One 12, e0175642 (2017).

16. Rasmussen, S. G. F. et al. Crystal structure of the β2 adrenergic receptor-Gs protein complex. Nature 477, 549–555 (2011).

17. Fan, Q. R. & Hendrickson, W. A. Structure of human follicle-stimulating hormone in complex with its receptor. Nature 433, 269–277 (2005).

18. Jiang, X. et al. Structure of follicle-stimulating hormone in complex with the entire ectodomain of its receptor. Proc. Natl. Acad. Sci. U. S. A. 109, 12491–12496 (2012).

19. Duan, J. et al. Structures of full-length glycoprotein hormone receptor signalling complexes. Nature (2021) doi:10.1038/s41586-021-03924-2.

20. Ulloa-Aguirre, A., Timossi, C., Damián-Matsumura, P. & Dias, J. A. Role of glycosylation in function of follicle-stimulating hormone. Endocrine 11, 205–215 (1999).

21. Caltabiano, G. et al. The specificity of binding of glycoprotein hormones to their receptors. Cell. Mol. Life Sci. 65, 2484–2492 (2008).

22. Moyle, W. R. et al. Co-evolution of ligand-receptor pairs. Nature 368, 251–255 (1994).

23. Lapthorn, A. J. et al. Crystal structure of human chorionic gonadotropin. Nature 369, 455–461 (1994).

24. Wu, H., Lustbader, J. W., Liu, Y., Canfield, R. E. & Hendrickson, W. A. Structure of human chorionic gonadotropin at 2.6 A resolution from MAD analysis of the selenomethionyl protein. Structure 2, 545–558 (1994).

25. Grossmann, M. et al. Substitution of the seat-belt region of the thyroid-stimulating hormone (TSH) beta-subunit with the corresponding regions of choriogonadotropin or follitropin confers luteotropic but not follitropic activity to chimeric TSH. J. Biol. Chem. 272, 15532–15540 (1997).

26. Dias, J. A., Zhang, Y. & Liu, X. Receptor binding and functional properties of chimeric human follitropin prepared by an exchange between a small hydrophilic intercysteine loop of human follitropin and human lutropin. J. Biol. Chem. 269, 25289–25294 (1994).

27. Chen, F., Wang, Y. & Puett, D. Role of the invariant aspartic acid 99 of human choriogonadotropin beta in receptor binding and biological activity. J. Biol. Chem. 266, 19357–19361 (1991).

28. Mueller, S., Jaeschke, H., Günther, R. & Paschke, R. The hinge region: an important receptor component for GPHR function. Trends Endocrinol. Metab. 21, 111–122 (2010).

29. Mizutori, Y., Chen, C.-R., McLachlan, S. M. & Rapoport, B. The Thyrotropin Receptor Hinge Region Is Not Simply a Scaffold for the Leucine-Rich Domain but Contributes to Ligand Binding and Signal Transduction. Molecular Endocrinology vol. 22 1171–1182 (2008).

30. Brüser, A. et al. The Activation Mechanism of Glycoprotein Hormone Receptors with Implications in the Cause and Therapy of Endocrine Diseases. J. Biol. Chem. 291, 508 (2016).

31. Costagliola, S. et al. Tyrosine sulfation is required for agonist recognition by glycoprotein hormone receptors. EMBO J. 21, 504 (2002).

32. Kosugi, S., Ban, T., Akamizu, T. & Kohn, L. D. Sitedirected mutagenesis of a portion of the extracellular domain of the rat thyrotropin receptor important in autoimmune thyroid disease and nonhomologous with gonadotropin receptors. Relationship of functional and immunogenic domains. J. Biol. Chem. 266, 19413–19418 (1991).

33. van Koppen, C. J. et al. Mechanism of action of a nanomolar potent, allosteric antagonist of the thyroidstimulating hormone receptor. Br. J. Pharmacol. 165, 2314–2324 (2012).

34. Van Sande, J. et al. In Chinese hamster ovary K1 cells dog and human thyrotropin receptors activate both the cyclic AMP and the phosphatidylinositol 4,5bisphosphate cascades in the presence of thyrotropin and the cyclic AMP cascade in its absence. Eur. J. Biochem. 229, 338–343 (1995).

35. Chen, C.-R., McLachlan, S. M. & Rapoport, B. A monoclonal antibody with thyrotropin (TSH) receptor inverse agonist and TSH antagonist activities binds to the receptor hinge region as well as to the leucine-rich domain. Endocrinology 150, 3401–3408 (2009).

36. Human monoclonal thyroid stimulating autoantibody. Lancet 362, 126–128 (2003).

37. Sanders, J. et al. Crystal structure of the TSH receptor in complex with a thyroid-stimulating autoantibody. Thyroid 17, 395–410 (2007).

38. Sanders, P. et al. Crystal structure of the TSH receptor (TSHR) bound to a blocking-type TSHR autoantibody. J. Mol. Endocrinol. 46, 81–99 (2011).

39. Evans, M. et al. Monoclonal autoantibodies to the TSH receptor, one with stimulating activity and one with blocking activity, obtained from the same blood sample. Clin. Endocrinol. 73, 404–412 (2010).

40. Schulze, A. et al. The intramolecular agonist is obligate for activation of glycoprotein hormone receptors. FASEB J. 34, 11243–11256 (2020).

41. Lomize, M. A., Pogozheva, I. D., Joo, H., Mosberg, H. L. & Lomize, A. L. OPM database and PPM web server: resources for positioning of proteins in membranes. Nucleic Acids Res. 40, D370–6 (2012).

42. Erbel, P. J. A., Haseley, S. R., Kamerling, J. P. & Vliegenthart, J. F. G. Studies on the relevance of the glycan at Asn-52 of the alpha-subunit of human chorionic gonadotropin in the alphabeta dimer. Biochem. J 364, 485–495 (2002).

43. Amr, S. et al. Activities of deglycosylated thyrotropin at the thyroid membrane receptor-adenylate cyclase system. J. Endocrinol. Invest. 8, 537–541 (2014).

44. Fares, F. A., Levi, F., Reznick, A. Z. & Kraiem, Z. Engineering a potential antagonist of human thyrotropin and thyroid-stimulating antibody. J. Biol. Chem. 276, 4543–4548 (2001).

45. Reily, C., Stewart, T. J., Renfrow, M. B. & Novak, J. Glycosylation in health and disease. Nat. Rev. Nephrol. 15, 346–366 (2019).

46. Feng, X., Müller, T., Mizrachi, D., Fanelli, F. & Segaloff, D. L. An intracellular loop (IL2) residue confers different basal constitutive activities to the human lutropin receptor and human thyrotropin receptor through structural communication between IL2 and helix 6, via helix 3. Endocrinology 149, 1705–1717 (2008).

47. Zhang, M. et al. Intrinsic differences in the response of the human lutropin receptor versus the human follitropin receptor to activating mutations. J. Biol. Chem. 282, 25527–25539 (2007).

48. Skiba, M. A. & Kruse, A. C. Autoantibodies as Endogenous Modulators of GPCR Signaling. Trends Pharmacol. Sci. 42, 135–150 (2021).

49. Asarnow, D., Palovcak, E. & Cheng, Y. UCSF pyem v0. 5. Zenodo https://doi.org/10.5281/zenodo 3576630, (2019).

50. Sanders, J. et al. Characteristics of a human monoclonal autoantibody to the thyrotropin receptor: sequence structure and function. Thyroid 14, 560–570 (2004).

51. Zheng, S. Q. et al. MotionCor2: anisotropic correction of beam-induced motion for improved cryo-electron microscopy. Nat. Methods 14, 331–332 (2017).

52. Punjani, A., Rubinstein, J. L., Fleet, D. J. & Brubaker, M. A. cryoSPARC: algorithms for rapid unsupervised cryo-EM structure determination. Nat. Methods 14, 290–296 (2017).

53. Punjani, A., Zhang, H. & Fleet, D. J. Non-uniform refinement: adaptive regularization improves singleparticle cryo-EM reconstruction. Nat. Methods 17, 1214–1221 (2020).

54. Scheres, S. H. W. A Bayesian view on cryo-EM structure determination. J. Mol. Biol. 415, 406–418 (2012).

55. Pettersen, E. F. et al. UCSF ChimeraX: Structure visualization for researchers, educators, and developers. Protein Sci. 30, 70–82 (2021).

56. Jumper, J. et al. Highly accurate protein structure prediction with AlphaFold. Nature 596, 583–589 (2021).

57. Emsley, P. & Cowtan, K. Coot: model-building tools for molecular graphics. Acta Crystallogr. D Biol. Crystallogr. 60, 2126–2132 (2004).

58. Zhuang, Y. et al. Mechanism of dopamine binding and allosteric modulation of the human D1 dopamine receptor. Cell Res. 31, 593–596 (2021).

59. Schüttelkopf, A. W. & van Aalten, D. M. F. PRODRG: a tool for high-throughput crystallography of protein– ligand complexes. Acta Crystallogr. D Biol. Crystallogr. 60, 1355–1363 (2004).

60. Croll, T. I. ISOLDE: a physically realistic environment for model building into low-resolution electrondensity maps. Acta Crystallogr D Struct Biol 74, 519–530 (2018).

61. Adams, P. D. et al. PHENIX: a comprehensive Python-based system for macromolecular structure solution. Acta Crystallogr. D Biol. Crystallogr. 66, 213–221 (2010).

62. Chen, V. B. et al. MolProbity: all-atom structure validation for macromolecular crystallography. Acta Crystallogr. D Biol. Crystallogr. 66, 12–21 (2010).

63. Bligh, E. G. A Rapid Method of Total Lipid Extraction and Purification. (1959).

64. MacLean, B. et al. Skyline: an open source document editor for creating and analyzing targeted proteomics experiments. Bioinformatics 26, 966–968 (2010).

65. Marty, M. T. et al. Bayesian deconvolution of mass and ion mobility spectra: from binary interactions to polydisperse ensembles. Anal. Chem. 87, 4370–4376 (2015).

