## Supplementary Information for "Autoantibody and hormone activation of the thyrotropin G protein-coupled receptor"

### Supplementary Figures

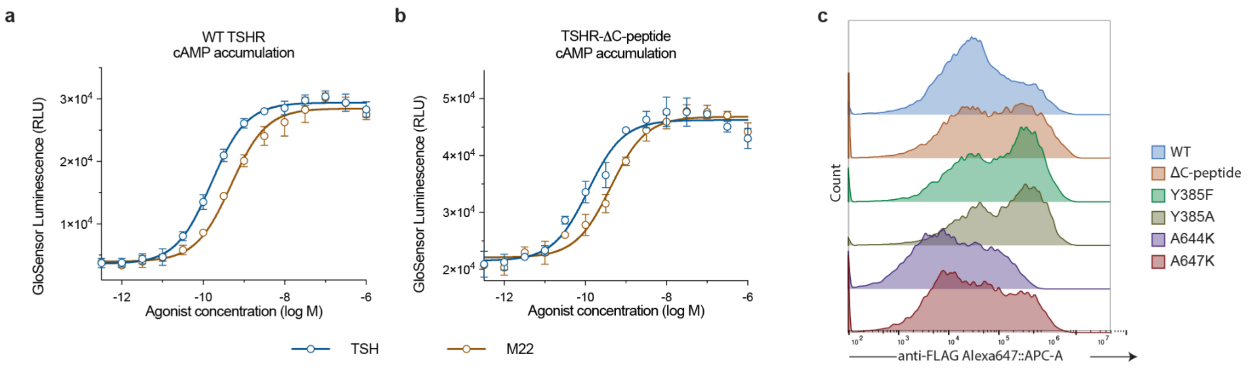


**Supplementary Figure 1 | Validation of TSHR mutants via signaling studies and cell surface staining.**
**a)** Glosensor cAMP accumulation assay for wild-type TSHR activated by TSH or M22 Fab. Signaling data points represent the global fit of grouped triplicate measurements at each concentration ± SD from 2 independent experiments. **b)** Glosensor cAMP accumulation assay for TSHR-ΔCpep activated by TSH or M22 Fab. **c)** Cell surface staining of wild-type TSHR and constructs used in signaling studies. Flag-tagged TSHR constructs were stained with Alexa647-labeled M1-FLAG antibody. Fluorescence from single cells was quantified by flow cytometry.

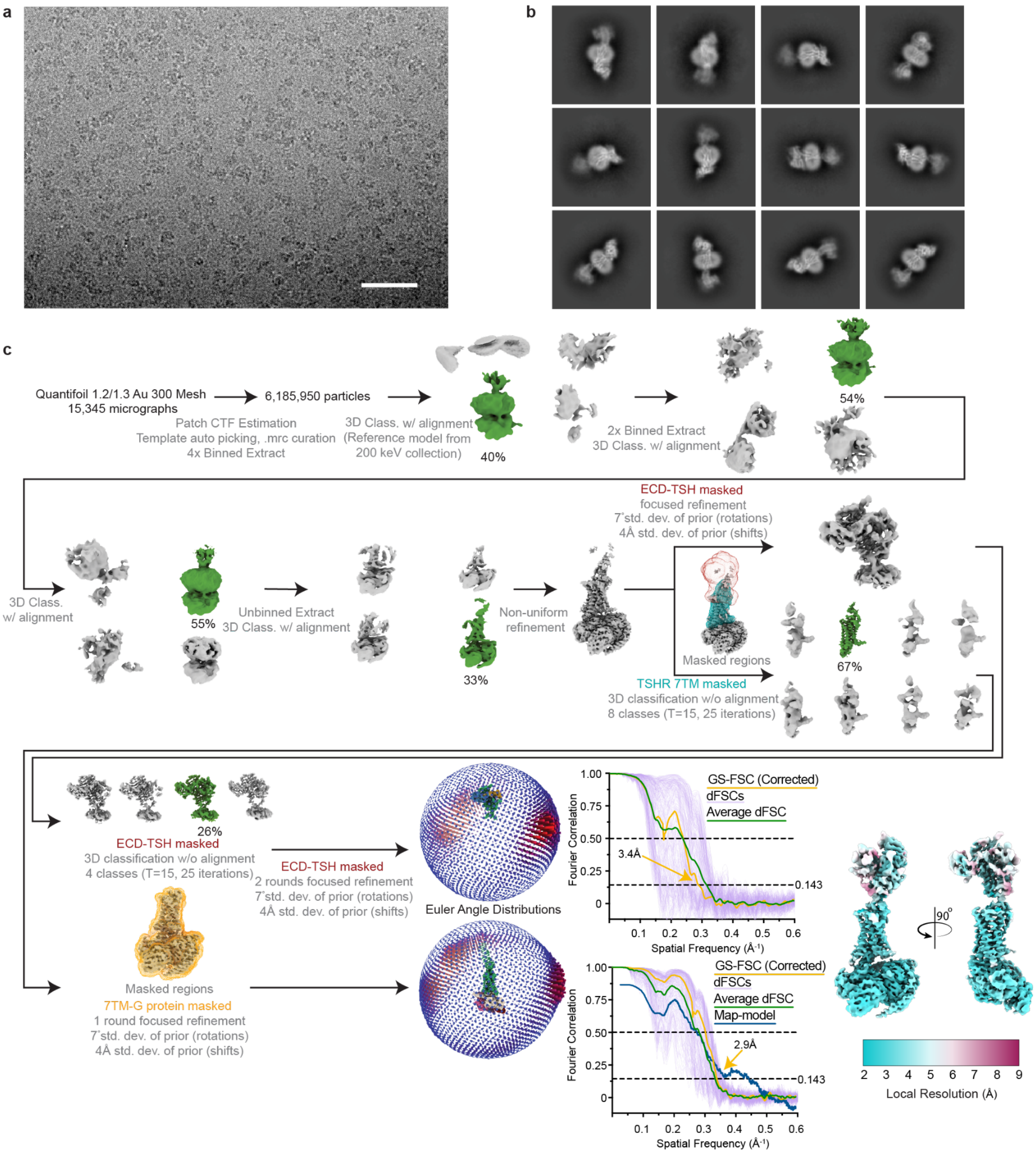


**Supplementary Figure 2 | Cryo-EM data processing for TSH-bound TSHR-$G_s$ complex**

**a)** Representative micrograph from data collection. Scale bar, 50 nm. **b)** Selected 2D class averages. **c)** Processing approach used for reconstruction of TSH-bound TSHR-$G_s$ complex. A local resolution map was calculated from cryoSPARC using masks from indicated local refinement, then visualized with the composite map in the same scale. A viewing distribution plot was generated using scripts from the pyEM software suite[49] and visualized in ChimeraX. GS-FSC and Directional FSC (dFSC, shown as purple lines) curves were generated in cryoSPARC and as previously described in Dang, S. *et al. Nature* 552, 426-429 (2017).

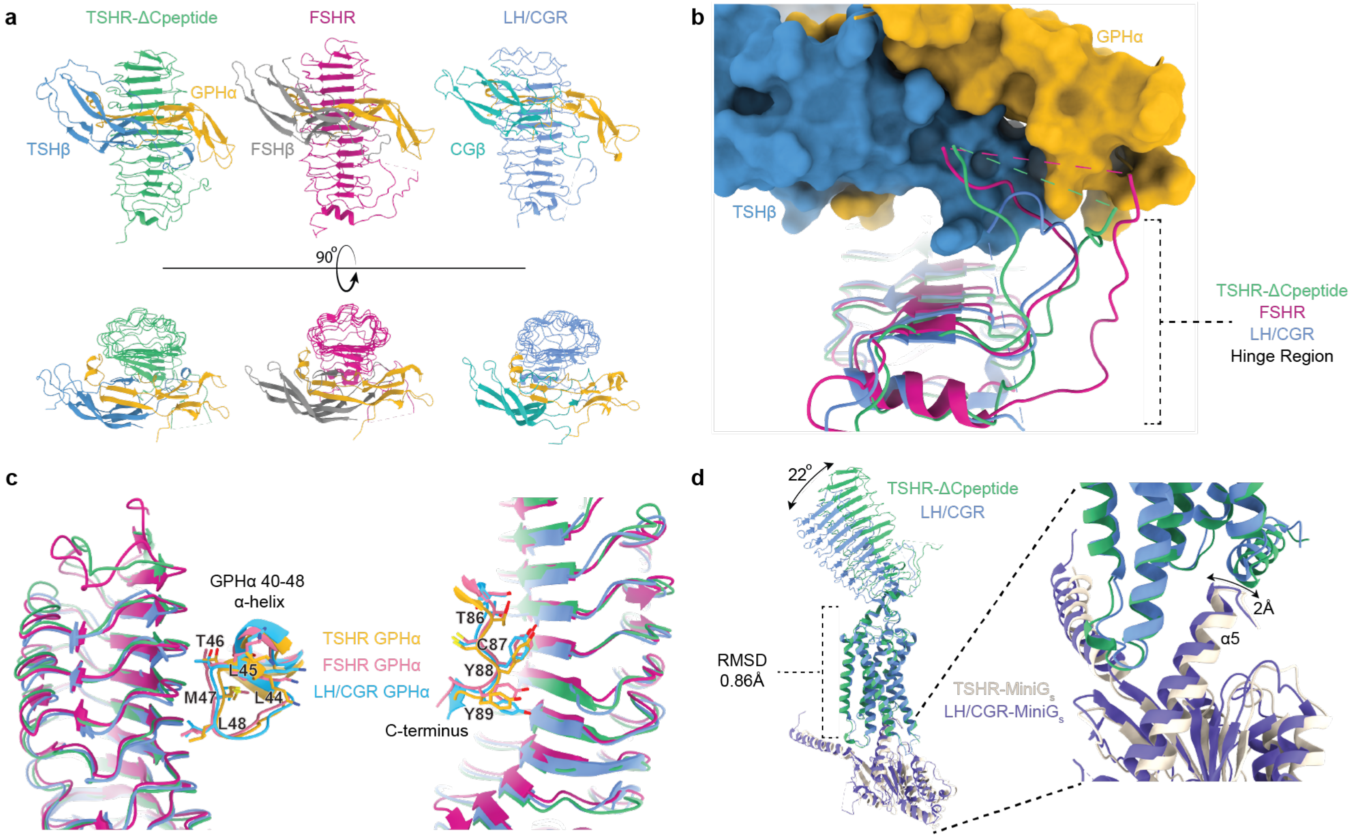


**Supplementary Figure 3 | Receptor:hormone interaction comparisons across the glycoprotein hormone receptor family.**
**a)** Comparison of TSHR-ΔCpeptide, FSHR, and LH/CGR ECD:hormone interactions shows the comparable concave binding-interface and lateral-ECD contacts made by GPHα and receptor-specific β subunits. **b)** Alignment of the GPHR ECDs reveals the conformational variability of the unstructured hinge region. Surface representation of TSH is shown. **c)** Alignment of GPHR ECDs highlighting GPHα contacts (labeled) on the lateral edges of GPHR ECDs that retain similar structural motifs and side chain orientations. **d)** Overall comparison of TSHR-ΔCpeptide and LH/CGR shows a high degree of structural similarity in transmembrane domain and, for CG-bound LH/CGR, a ~22° rotation of the ECD towards the membrane. Overall $G_s$ interactions are highly similar between TSHR-ΔCpeptide and LH/CGR (TM6/7 hidden).

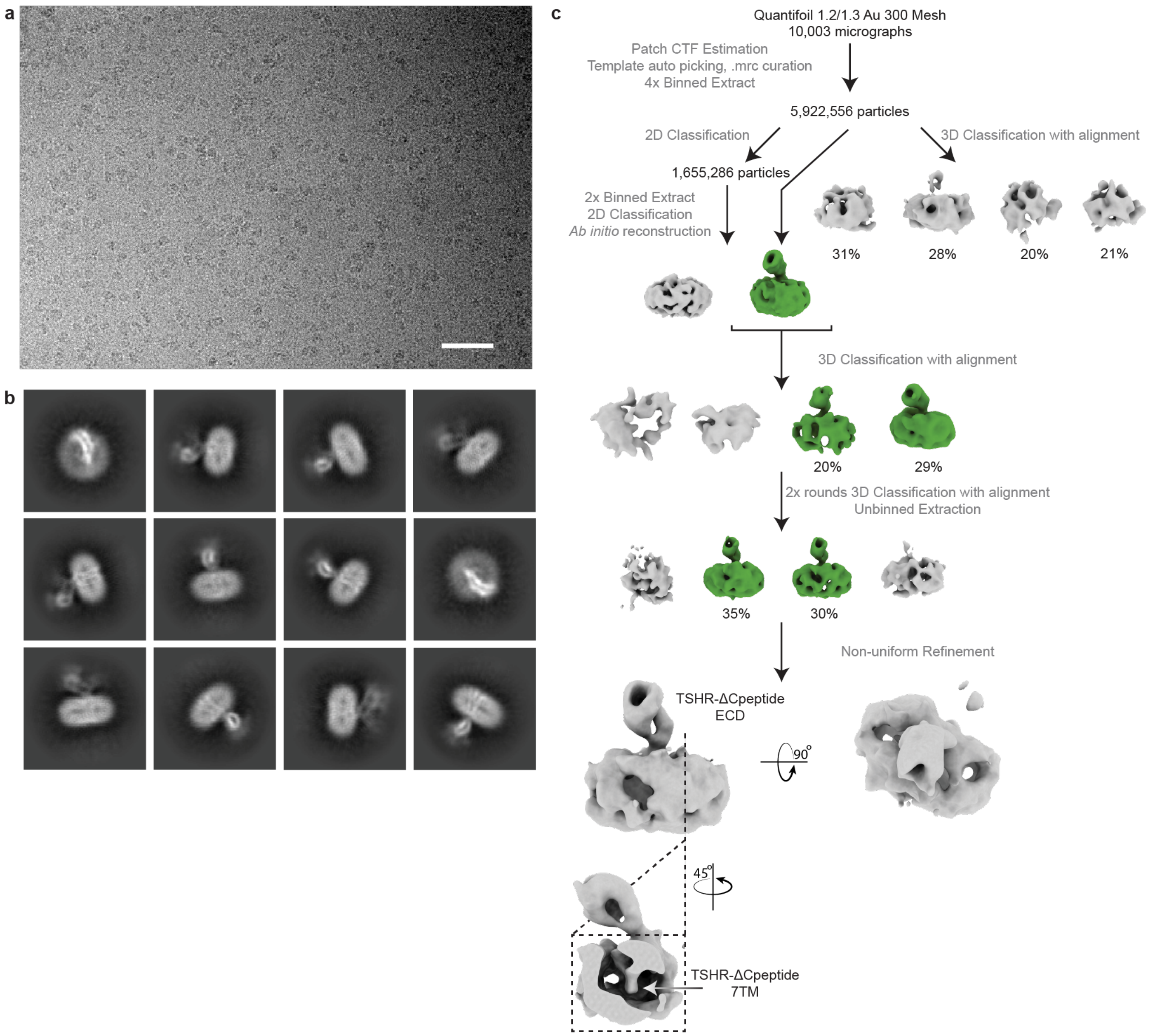


**Supplementary Figure 4 | Cryo-EM data processing for Org 274179-0 bound TSHR. a)** Representative micrograph from data collection. Scale bar, 50 nm. **b)** Selected 2D class averages generated from curated particles. **c)** Processing approach used for low resolution reconstruction. Despite starting with a similar or larger sized dataset as for other TSHR samples, the TSHR-Org 274179-0 complex does not yield high resolution reconstruction of the 7TM domain. A low-resolution reconstruction of the TSHR ECD was observed. This suggests potential flexibility between the TSHR 7TM domain and the ECD.

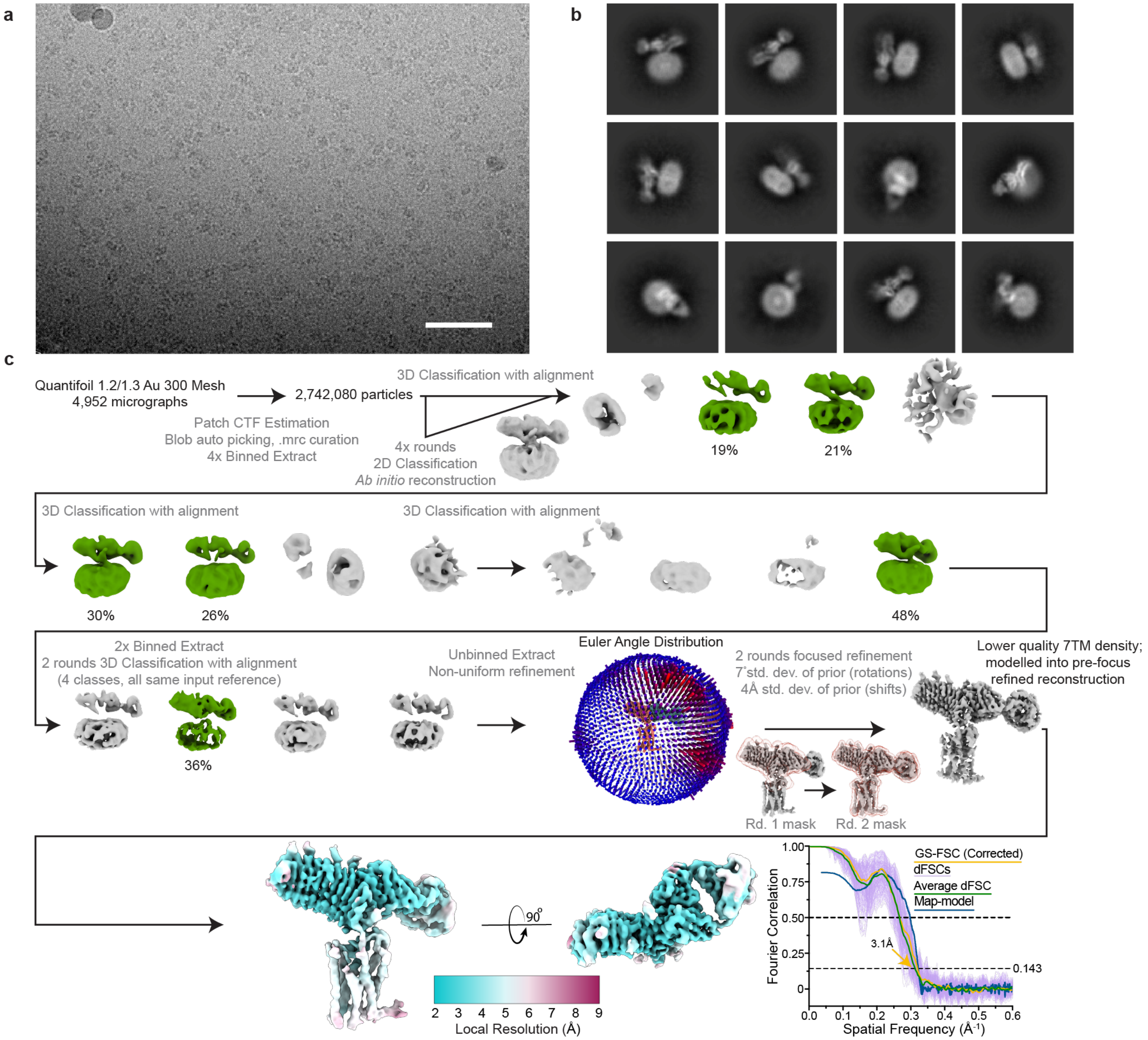


**Supplementary Figure 5 | Cryo EM data processing for the CS-17 bound TSHR:Org 274179-0 complex. a)** Representative micrograph from data collection. Scale bar, 50 nm. **b)** Selected 2D class averages generated from the final reconstruction. **c)** Processing approach used for reconstruction of the complex. A viewing distribution plot was generated using scripts from the pyEM software suite and visualized in ChimeraX. Local resolution map generated from non-uniform refinement mask in cryoSPARC. GS-FSC and dFSC curves were generated in cryoSPARC and as previously described in Dang, S. *et al. Nature* 552, 426-429 (2017).

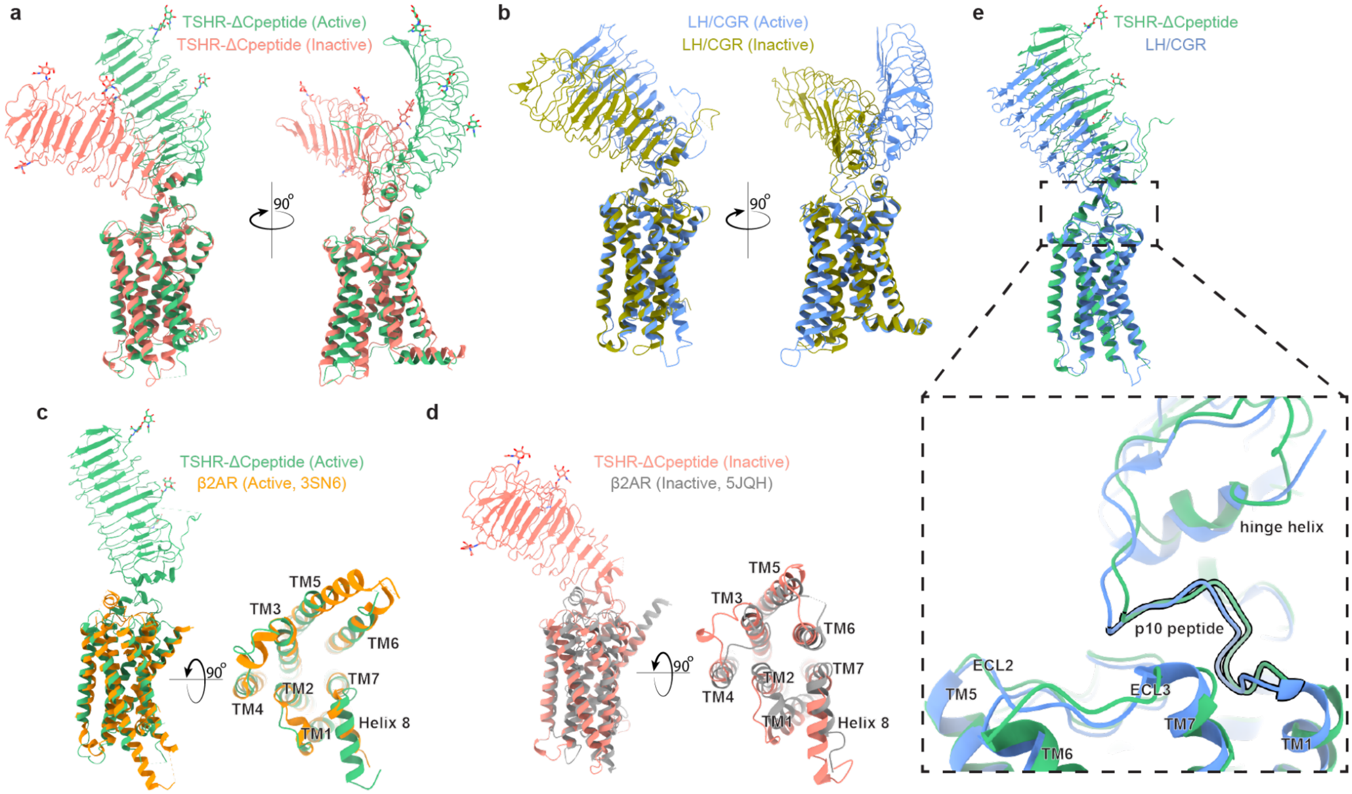


**Supplementary Figure 6 | Comparison of TSHR activation with other GPCRs.**
**a)** Ribbon diagram of TSHR in active and inactive conformations. **b)** Ribbon diagram of LH/CGR in active and inactive conformations. A similar reorientation of the ECD is shared between TSHR and LH/CGR upon activation. **c)** Comparison of active TSHR to active β2-adrenoceptor (β2AR). **d)** Comparison of inactive TSHR to inactive β2AR. **e)** Comparison of active conformations of TSHR and LH/CGR reveals similar overall structures of the 7TM domain and the p10 peptide.

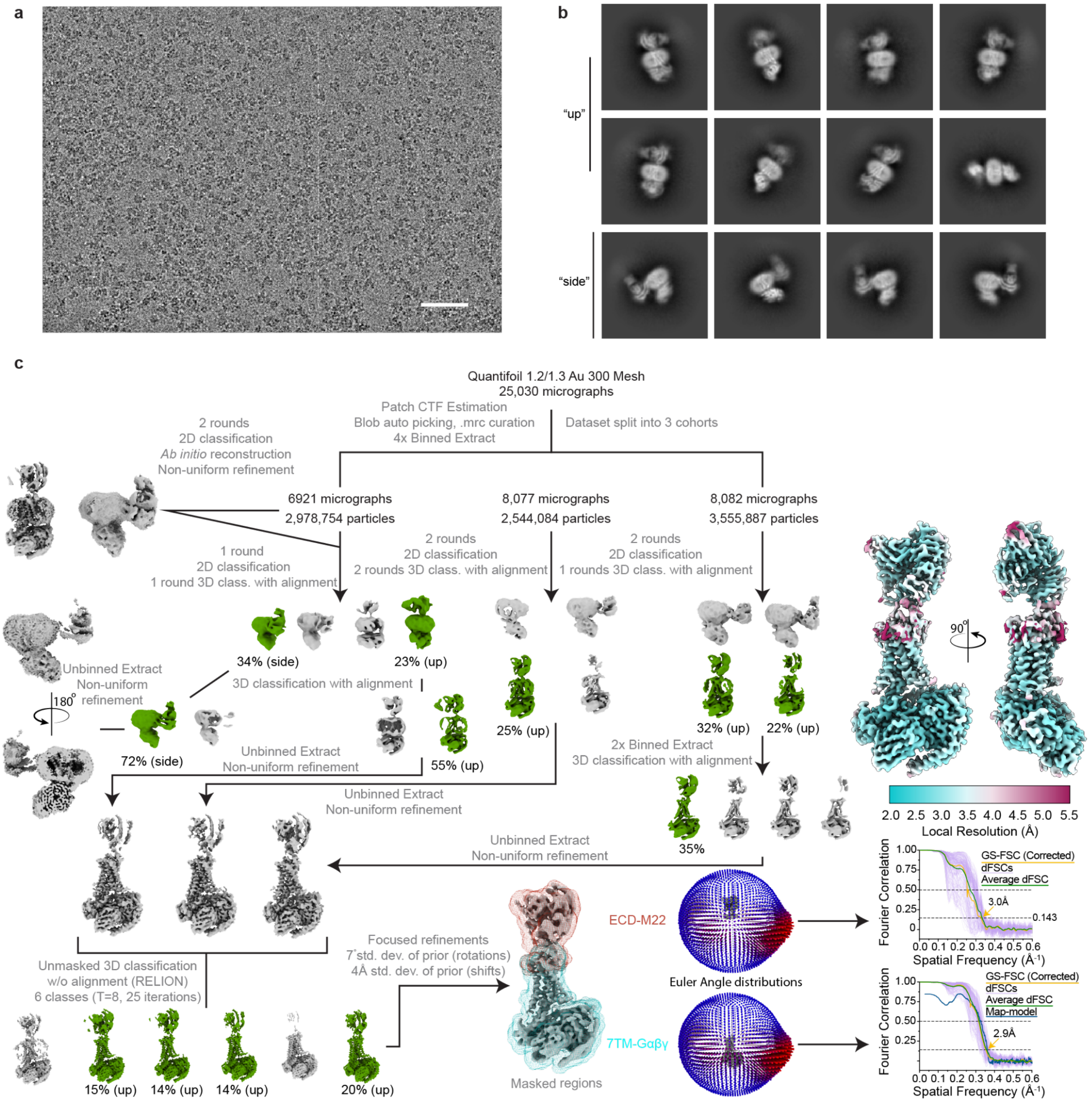


**Supplementary Figure 7 | Cryo-EM data processing for TSH-bound TSHR-G$_s$ complex.**
**a)** Representative micrograph from data collection. Scale bar, 50 nm. **b)** Selected 2D class averages from final reconstruction. **c)** Processing approach used for reconstruction of M22-bound TSHR-G$_s$ complex. A local resolution map was calculated from cryoSPARC using masks from indicated local refinement, then visualized with the composite map in the same scale. A viewing distribution plot was generated using scripts from the pyEM software suite and visualized in ChimeraX. GS-FSC and dFSC curves were generated in cryoSPARC and as previously described in Dang, S. *et al. Nature* 552, 426-429 (2017).

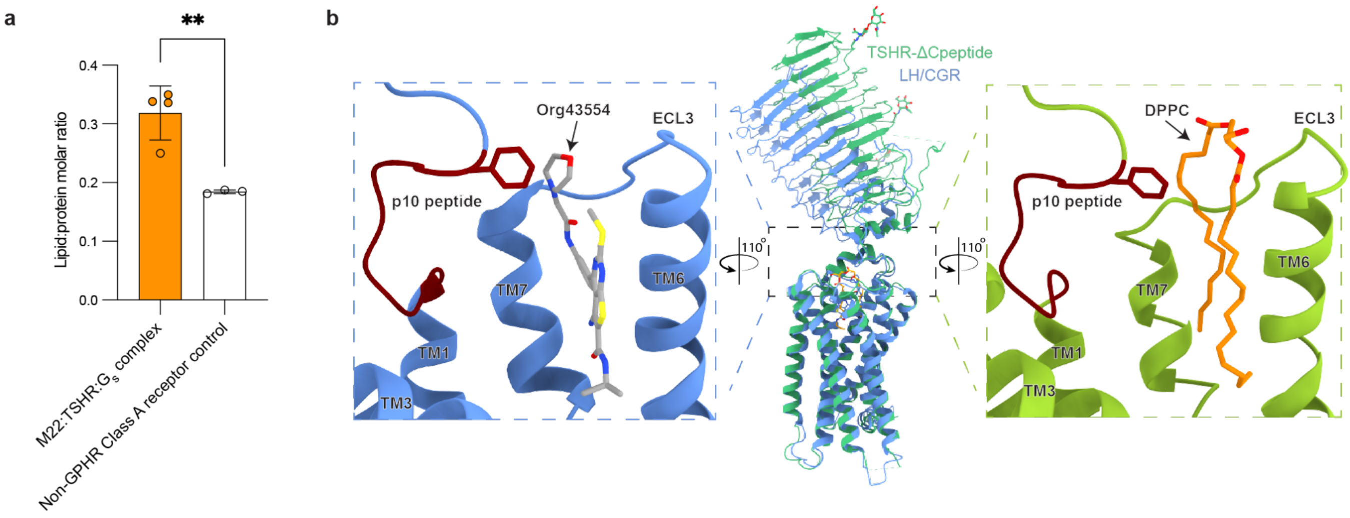


**Supplementary Figure 8 | Lipid binding site in TSHR.**

**a)** Lipid:protein molar ratio comparison between the M22:TSHR:$G_s$ complex and a non-GPHR Class A receptor control. Data points represent individual measurements of the ratio of pmol of lipid DPPC per pmol of protein. $^{**}P = 0.0088$; Unpaired two-tailed t test was used to calculate statistical differences in lipid:protein molar ratios. **b)** Comparison of the TSHR-ΔCpeptide and LHCGR transmembrane pockets highlights the similarity in the DPPC and the allosteric agonist Org43554 binding sites, further suggestive of the mechanistic importance of this endogenous lipid in TSHR. TM4/5 and ECL2 not shown.

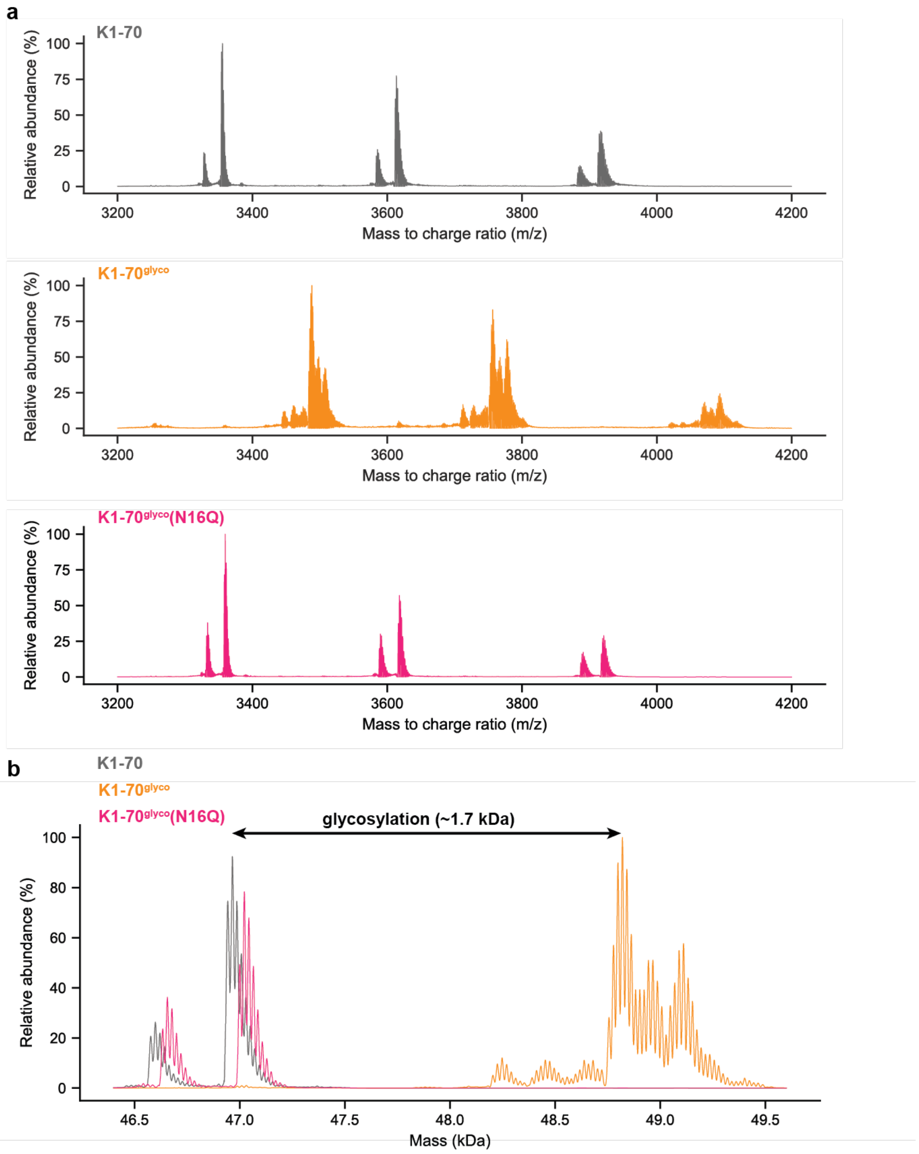


**Supplementary Figure 9 | Glycosylation of engineered K1-70$^{glyco}$ construct. a)** Native mass spectrum (nMS) of K1-70, K1-70$^{glyco}$, and K1-70$^{glyco}$(N16Q) Fab fragments. **b)** Accurate mass assignment of Fab fragments. nMS demonstrates ~1.7 kDa increased mass for K1-70$^{glyco}$ and more heterogeneity consistent with N-linked glycosylation. The smaller mass and increased homogeneity of the K1-70$^{glyco}$(N16Q) construct further supports glycosylation at the engineered N16 position.

**Supplemental Table 1. Cryo-EM data collection, refinement and validation statistics**

| | TSH-bound TSHR-$G_s$<br><br>(EMDB-xxxx)<br>(PDB xxxx) | Org 2274179-0-bound TSHR<br><br>(EMDB-xxxx)<br>(PDB xxxx) | CS-17 Fab/ Org 2274179-0-bound TSHR<br>(EMDB-xxxx)<br>(PDB xxxx) | M22-bound TSHR-$G_s$<br><br>(EMDB-xxxx)<br>(PDB xxxx) |
|---|---|---|---|---|
| **Data collection and processing** | | | | |
| Magnification | 130,000 | 105,000 | 130,000 | 81,000 |
| Voltage (kV) | 300 | 300 | 300 | 300 |
| Electron exposure (e–/Å$^2$) | 77 | 50 | 77 | 60 |
| Defocus range (μm) | -0.8 to -2.2 | -0.8 to -2.4 | -0.8 to -2.2 | -0.8 to -2.0 |
| Pixel size (Å) | 0.662 (physical) | 0.85 (physical) | 0.644 (physical) | 0.844 (physical) |
| Symmetry imposed | *C1* | *C1* | *C1* | *C1* |
| Initial particle images (no.) | 6,185,950 | 5,922,556 | 2,742,080 | 9,078,725 |
| Final particle images (no.) | 80,483 | 357,869 | 41,054 | 244,973 |
| Map resolution (Å)<br>(masked) | 2.9 (7TM-miniG$\alpha_s\beta\gamma$)<br>3.4 (ECD-TSH) | 5.3 | 3.1 | 2.9 (7TM-miniG$\alpha_s\beta\gamma$)<br>3.0 (ECD-M22) |
| FSC threshold | 0.143 | 0.143 | 0.143 | 0.143 |
| Map resolution range (Å) | 2.9 – 9.0 | 5.3-50 | 3.1 – 7.0 | 2.9 - 8.0 |
| **Refinement** | | | | |
| Initial model used<br>(PDB code) | AlphaFold (TSH)<br>M22-bound TSHR-$G_s$ structure for remaining components | | AlphaFold (TSHR)<br>AlphaFold (CS-17) | AlphaFold (TSHR),<br>7LJC (G protein)<br>3SN6 (Nb35)<br>3G04 (M22) |
| Model resolution (Å)<br>(unmasked/masked) | 3.9/3.6 | | 3.5/3.3 | 3.2/3.1 |
| FSC threshold | 0.5 | | 0.5 | 0.5 |
| Model resolution range (Å) | 3.6-50 | | 3.5-50 | 3.2-50 |
| Map sharpening *B* factor (Å$^2$) | -102 (ECD),<br>-83 (7TM/MiniG$\alpha_s\beta\gamma$) | | -91 | -90 (ECD)<br>-80<br>(7TM/MiniG$\alpha_s\beta\gamma$) |
| Model composition | | | | |
| Non-hydrogen atoms | 11914 | | 7827 | 13405 |
| Protein residues | 1520 | | 995 | 1723 |
| Ligands | NAG: 8 | | NAG: 5 | NAG: 3 |
| *B* factors (Å$^2$) | | | | |
| Protein | 103.1 | | 93.5 | 89.9 |
| Ligand | 120.6 | | 49.4 | 45.6 |
| R.m.s. deviations | | | | |
| Bond lengths (Å) | 0.004 | | 0.006 | 0.006 |
| Bond angles (°) | 0.849 | | 1.221 | 0.675 |
| Validation | | | | |
| MolProbity score | 1.54 | | 1.53 | 1.58 |
| Clashscore | 4.23 | | 5 | 5.62 |
| Poor rotamers (%) | 0 | | 0.91 | 0.07 |
| CaBLAM outliers (%) | 1.49 | | 1.63 | 1.18 |
| Ramachandran plot | | | | |
| Favored (%) | 95.19 | | 96.05 | 96.05 |
| Allowed (%) | 4.81 | | 3.95 | 3.95 |
| Disallowed (%) | 0 | | 0 | 0 |

| TSHR WT/ΔC peptide cAMP accumulation | | | |
|---|---|---|---|
| TSHR Construct | Ligand | cAMP $EC_{50}$ | cAMP $E_{max}$ |
| WT | TSH | 2.0E-10 ± 0.1 (6) | 100 ± 5.5 (6) |
| | M22 | 8.3E-10 ± 0.04 (6) | 94.6 ± 4.8 (6) |
| ΔC Peptide | TSH | 1.5E-10 ± 0.1 (6) | 84.5 ± 8.9 (6) |
| | M22 | 5.2E-10 ± 0.09 (6) | 79 ± 12.1 (6) |

| Sulfotyrosine mutant cAMP accumulation | | | |
|---|---|---|---|
| TSHR Construct | Ligand | cAMP $EC_{50}$ | cAMP $E_{max}$ |
| WT | TSH | 2.2E-10 ± 0.12 (6) | 100 ± 3.9 (6) |
| | M22 | 4.8E-10 ± 0.28 (6) | 89.4 ± 0.1 (6) |
| TSHR Y385F | TSH | 6.2E-10 ± 0.02 (6) | 98.6 ± 2.6 (6) |
| | M22 | 1.4E-09 ± 0.3 (6) | 85.3 ± 12.3 (6) |
| TSHR Y385A | TSH | 8.7E-10 ± 0.03 (6) | 116.8 ± 29 (6) |
| | M22 | 8.7E-10 ± 0.03 (6) | 80.5 ± 2.5 (6) |

| Lipid site mutant cAMP accumulation | | | |
|---|---|---|---|
| TSHR Construct | Ligand | cAMP $EC_{50}$ | cAMP $E_{max}$ |
| WT | M22 | 2.9E-10 ± 0.04 (16) | 100 ± 1.9 (16) |
| TSHR A644K | M22 | 1.5E-8 ± 0.04 (16) | 54.7 ± 3.1 (16) |
| TSHR A647K | M22 | 2.2E-08 ± 0.08 (16) | 58.5 ± 3.6 (16) |

| K1-$70^{glyco}$ cAMP accumulation | | | |
|---|---|---|---|
| TSHR Construct | Ligand | cAMP $EC_{50}$ | cAMP $E_{max}$ |
| WT | M22 | 6.8E-10 ± 0.004 (18) | 100 ± 2.5 (18) |
| | K1-70 | 5.9E-09 ± 0.02 (18) | 35.3 ± 1.9 (18) |
| | K1-$70^{glyco}$ | 2.2E-09 ± 0.04 (18) | 122.8 ± 8.6 (18) |
| | K1-70 (N16Q) | 1.1E-08 ± 0.008 (18) | 49.9 ± 0.9 (18) |

**Supplementary Table 2 | Summary of TSHR signaling studies.** Values are expressed as mean $EC_{50}$ or mean $E_{max}$ ± s.e.m. from (n) technical replicates.
